# Differential responses and recovery dynamics of HSPC populations following *Plasmodium chabaudi* infection

**DOI:** 10.1101/2024.09.25.614952

**Authors:** Federica Bruno, Christiana Georgiou, Deirdre Cunningham, Samantha Atkinson, Lucy Bett, Marine Secchi, Sara Gonzalez Anton, Flora Birch, Jean Langhorne, Cristina Lo Celso

## Abstract

Severe infections such as malaria are on the rise worldwide, driven by both climate change and increasing drug-resistance. It is therefore paramount that we better understand how the host’s organism responds to severe infection. Hematopoiesis is particularly of interest in this context because hematopoietic stem and progenitor cells (HSPCs) maintain the turnover of all blood cells, including all immune cells. Severe infections have been widely acknowledged to affecting HSPCs, however this disruption has been mainly studied during the acute phase, and the process and level of HSPC recovery remains understudied. Using a self-resolving model of natural rodent malaria, infection by *Plasmodium chabaudi*, here we systematically assess HSPCs’ acute response and recovery upon pathogen clearance. We demonstrate that during the acute phase of infection the most quiescent and functional stem cells are depleted, multipotent progenitor compartments are drastically enlarged, and oligopotent progenitors virtually disappear, underpinned by dramatic, population-specific and sometimes unexpected changes in proliferation rates. HSPC populations return to homeostatic size and proliferation rate again through specific patterns of recovery. Overall, our data demonstrate that HSPC populations adopt different responses to cope with severe infection and suggest that the ability to adjust proliferative capacity becomes more restricted as differentiation progresses.

## INTRODUCTION

Mature blood and immune cells are continuously replenished through a dynamic process known as hematopoiesis. During homeostasis, this process is tightly regulated and characterized by a hierarchical cellular differentiation system so that the high turnover of blood cells is maintained lifelong. At the top of the hematopoietic system, a pool of hematopoietic stem cells (HSCs) is able to generate all differentiated blood cells types by giving rise to several different progenitors thanks to their self-renewal and multilineage differentiation capacity^1^. It is now clear that while most HSCs usually are in a quiescent state, the massive daily turnover of blood cells is sustained by a highly proliferative pool of progenitors, from multipotent progenitors (MPPs), direct progeny of HSCs, to more mature, oligopotent progenitors^2^. A wealth of studies has characterized hematopoietic stem and progenitor cell (HSPC) populations^3^. Of note, self-renewal and multilineage potential have been shown to decrease alongside the lineage commitment process. While MPP populations are lineage biased but retain multipotentiality, common myeloid progenitors (CMPs), granulocyte-macrophage progenitors (GMPs), megakaryocyte-erythrocyte progenitors (MEPs), and common lymphoid progenitors (CLPs) are committed to the lineages indicated by their names^4^. Overall, decades of research have shown that these populations differ in terms of proliferation kinetics, homeostatic functions, maintenance mechanisms and lifespan, highlighting the high heterogeneity present within the hematopoietic system and even within the HSC pool itself.

During hematopoietic stress, the dynamics of this tightly regulated but flexible system change, leading to disrupted hematopoiesis. In response to several model of infections, HSCs are forced out of quiescence in order to cope with the increased demand for immune cells, which are consumed during the fight against the pathogen ^5–7^. Collectively HSPCs have been shown to sense and respond to cytokines and pro-inflammatory stimuli, resulting in several phenomena such as increased proliferation and differentiation towards the myeloid lineage, leading to emergency myelopoiesis, as well as increased mobilization outside of the bone marrow (BM) ^8,9^. Many studies reported that forced entry into the cell cycle resulted in replication stress within HSCs, leading to their loss of functionality, demonstrated by a decrease in repopulation potential in transplantation assays^10,11^. Following transient infections, HSCs have been shown to return to a dormant state, while continuous stimulation to proliferate in response to chronic inflammation consequently led to accelerated stem cell exhaustion and long-term functional impairment ^12–17^. In the latter, the ability of some HSCs to return to quiescence has been proposed to be critical to retain some function ^11^.

Malaria is a severe and life-threatening disease and still a leading cause of mortality. Of the more than 240,000 million cases of malaria in humans per year, most are resolved; however, there are more than 600,000 deaths per year, 90% of which are caused by *P. falciparum* in children in sub-Saharan Africa^18^. Following entrance into the host through the skin, *Plasmodium* undergoes a first amplification step in the liver, and once matured it is released into the bloodstream where it finally invades red blood cells (RBCs) leading to anemia and altered erythropoiesis. Malaria has been reported to compromise the immune system both in the short-term, leading to complications such as cerebral malaria, acute respiratory distress and kidney dysfunction, and in the long-term, resulting in severe anemia and loss of BM cellularity ^19–22^. The pathophysiology of malaria is centered around acute systemic inflammation, which results in pro-inflammatory cytokines production, lymphocyte activation and vessel congestion by adhesion of infected RBCs to endothelium ^19,23^. Recently, parasites have also been documented to circulate, reside and infect cells in the BM, suggesting the BM microenvironment as a potential *Plasmodium* reservoir ^19,22^. Acute *Plasmodium* infection has been proven to alter hematopoiesis, for instance a *Plasmodium*-induced IL-7Rα^+^c-Kit^high^ myeloid-primed progenitor population was reported to aid in the clearance of infected RBCs^24^. Analyses of primitive HSPC populations in mice infected by *Plasmodium berghei* highlighted emergency myelopoiesis at the expense of erythroid and lymphoid lineages, loss of functional HSCs and a role for the BM microenvironment in mediating such damage and as a possible therapeutic target to preserve HSC function during the acute phase of the infection ^7,25^. The extent and dynamics of HSPC recovery following malaria and other severe yet non-fatal infections have remained understudied, despite holding clues on potential strategies to alleviate both short and long-term consequences of infection. These are particularly important for malaria because the severe forms of this disease tend to occur in children. Generally, experimental strong inflammatory stresses have been linked to loss of HSC function, HSC ageing, and selection of HSC clones carrying mutations linked to clonal hematopoiesis and, eventually, malignant transformation ^10,15^. However, less is known about recovery from physiological severe infections.

While *P. berghei* is an excellent model of severe and cerebral malaria, it is fatal to mice unless treated with drugs that are themselves likely to affect hematopoiesis. Here we employ a different model of rodent malaria, *Plasmodium chabaudi chabaudi*, from which mice can recover without any pharmacological intervention. Through a time course analysis, we systematically characterize the effects of *P. chabaudi* on primitive HSPCs (HSCs and all MPP populations) and downstream oligopotent progenitor (CMP, GMP, MEP) populations in the BM throughout the acute and recovery stages. Immunophenotypic and EdU incorporation assays allow us to demonstrate that while eventually hematopoiesis returns to homeostatic conditions, HSPC populations are heterogenous in their response and recovery dynamics. These findings have important implications for better harnessing hematopoiesis to enable both infection resolution and healthy ageing.

## METHODS

### ANIMALS

All animal work, including animal care and experimental procedures, was carried out at Imperial College London and at the Francis Crick Institute in accordance with the current UK Home Office regulations (ASPA 1986) under license number PP9504146. C57BL/6 wild type (WT) mice were bred and housed at the Francis Crick Institute. Female mice aged 6-10 weeks were used for all experimental procedures as *P. chabaudi* infection has been reported to cause more severe complications in male C57BL/6 mice that compromise welfare.

### *P. CHABAUDI* EXPERIMENTAL MODEL

A cloned line of *Plasmodium chabaudi chabaudi* (AS) was used to initiate infections by intraperitoneal injection (*i.p.*) of 1 x 10^5^ infected RBCs collected from infected donor mice as previously described^26^. Blood-stage parasitemia was monitored by light microscopy assessment of Giemsa-stained (48900-500ML-F, Sigma-Aldrich) thin blood films generated from small drops of bloods collected from the tail veins on the following days: 4, 7, 8, 10, 11, 14, 17, 21, 24, 29, 35, 40, 45, 50, 55, 60 post infection (p.i.). The average parasitemia was measured as a percentage of at least 1000 RBCs counted per slide. Mice were sacrificed at the time points of interest, up to two months p.i..

### BLOOD CELL COUNTS ANALYSIS

Blood was collected via cardiac puncture at the time of sacrifice and transferred in ethylenediaminetetraacetic acid (EDTA)-coated tubes. To determine the total count of white blood cells (WBCs), RBCs and platelets, the blood was first diluted (1 in 5) in phosphate-buffered saline (PBS) and then run through the automatic hematology analyzer Sysmex XP-100.

### EdU INCORPORATION ASSAY

For cell populations’ proliferation analysis, 5-ethynyl-2’-deoxyuridine (EdU) was administered 2.5 hours prior the culling via intravenous injection (1mg/mouse). EdU, a thymidine analogue, is incorporated in newly synthesized DNA thus labelling the cells in the S-phase of the cell cycle. EdU uptake was measured by flow cytometry (see next section).

### FLOW CYTOMETRY

For the phenotypic analysis of HSPCs during infection, femurs, tibias and hips were harvested at day 8, day 11, day 15, day 24, day 29 and day 60 p.i. from age-matched healthy control and infected mice. Bones were crushed in FACS buffer (PBS, 2% fetal bovine serum (FBS)) and obtained cells were filtered through 40-µm strainer depleted of RBCs and stained with relevant monoclonal antibodies. For a comprehensive summary of antibody concentrations, clones and manufacturing companies consult Table S1. Dead cells were excluded by staining the cells with fixable viability stain 510 (BD Horizon).

For EdU detection, the Click-iT™ EdU Pacific Blue™ Kit (Invitrogen) was employed according to manufacturer’s instructions. Briefly, single cell suspensions were stained with the relevant antibodies, then fixed with 4% paraformaldehyde (PFA) and permeabilized, followed by incubation with the Click-iT™ detection cocktail.

To determine the absolute cell number in the populations of interest, Sysmex XP-100 or Calibrite beads (BD Biosciences) were used. For the former, the number of WBCs in two legs was multiplied by the frequency of single cells obtained via flow cytometry analysis. For the latter, Calibrite beads were added to the suspension prior to running the sample through the flow cytometer. Absolute numbers were then calculated by multiplying the beads added (100000) to the cell count of each population and divided by the number of beads detected by the machine.

Relevant controls, such as fluorescent minus one (FMOs) and single-color controls were included for compensation and gating purposes. All samples were analyzed on an LSR Fortessa (BD Biosciences) III, and the software FlowJo (Tree Star) was used to analyze the resulting data.

### STATISTICAL ANALYSIS

Raw data were tabulated in Microsoft excel, while graphs and statistical tests were created and performed via GraphPad Prism. Data are expressed as mean ± standard error of the mean (s.e.m.). Where stated, absolute cell numbers were normalized by dividing each value by the average of the corresponding control value for that day. For multiple comparisons, unpaired two tailed *t*-test with Holm–Šidák correction was used^27^. Differences were considered significant where (*) p < 0.05, (**) p < 0.01, (***) p < 0.001, (****) p < 0.0001. Detailed statistical information can be found in each figure caption. For all experiments, *n* refers to the total number of replicates at each timepoint, pooled from three independent experiments.

## RESULTS

### *Plasmodium chabaudi* infection is an effective model to study perturbed hematopoiesis and its recovery

To investigate the response of hematopoietic cells during acute infection and following resolution, C57BL/6 wild type mice were either mock injected or infected with *Plasmodium chabaudi* and their peripheral blood (PB) and BM were analyzed at different timepoints following infection (Figure 1A). The *P. chabaudi* blood-stage, acute infection in C57BL/6 mice was characterized by peak parasitemia at day 11 p.i., which was then resolved to subpatent levels by day 24 p.i. (Figure 1B), apart from a small patent recrudescence at day 29 p.i., consistent with previous observations^28–30^. Using parasitemia levels as a guideline, we defined different stages of infection and selected appropriate time points to investigate specific responses associated with different phases of infection. Namely, PB and BM samples from control and infected mice were analyzed at day 8 (early response, pre parasitemia peak), at day 11 and 15 (acute phase, peak and post peak of parasitemia respectively) and day 24, 29 and 60 (recovery phase, post-peak, very low to undetectable parasitemia) p.i. (Figure 1A-B).

**Figure 1.**
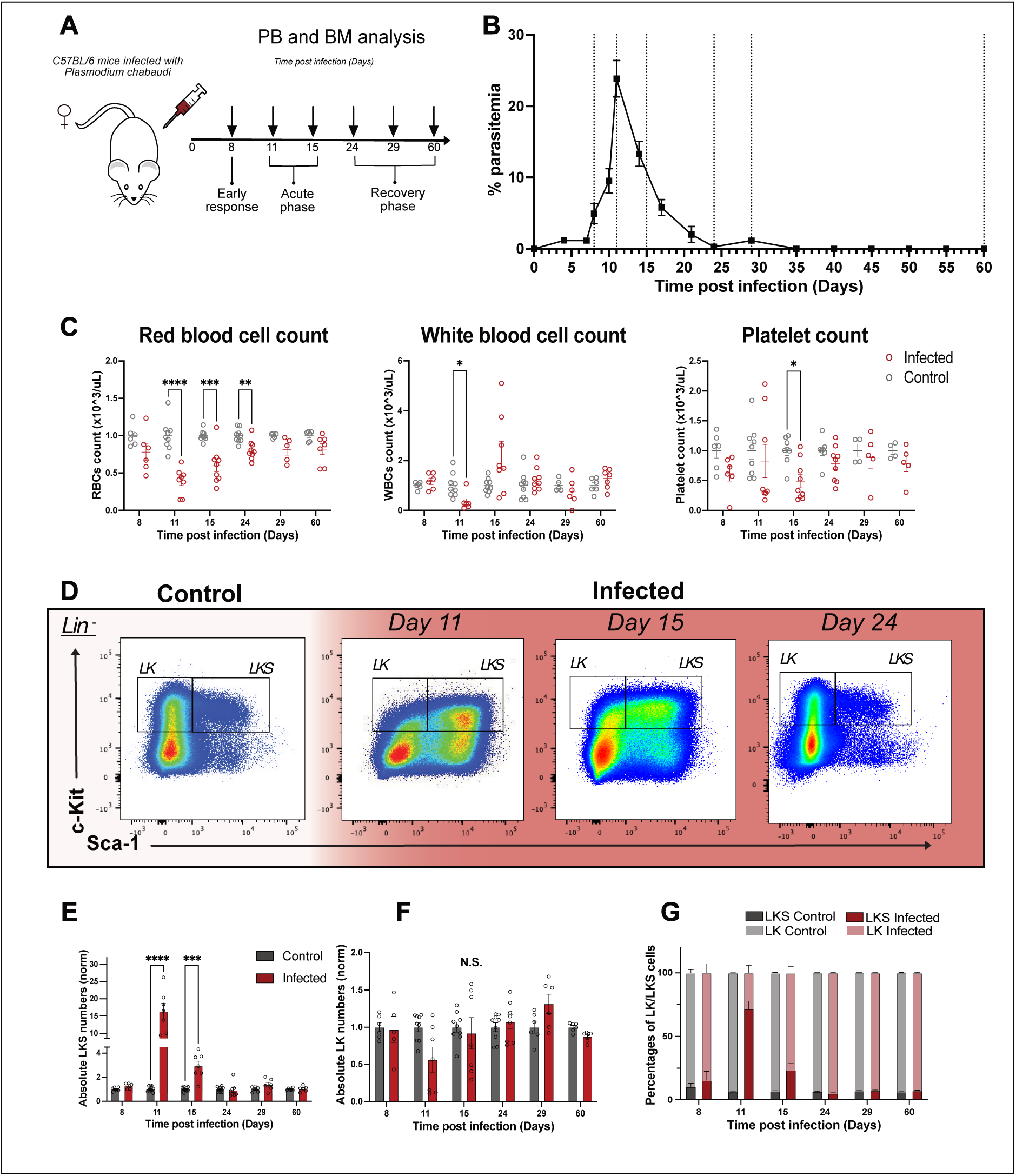
*P. chabaudi* is an effective model to study perturbed hematopoiesis and its recovery. **A**. Experimental set up. On day 0, C57BL/6 mice were injected with *P. chabaudi* infected RBCs and PB and BM analyses were conducted at days 8, 11, 15, 24, 29 and 60 p.i.. Note time is not to scale. **B.** Percentages of infected RBCs in peripheral blood over the course of the infection. Dotted lines highlight days when PB and BM were analyzed. **C.** Peripheral blood cell counts in control and infected mice during the infection, including RBC, WBC and platelet counts. **D.** Representative flow cytometry pseudo-color density plots showing LKS and LK populations in control and infected mice at day 11, 15 and 60 p.i. **E-F.** Normalized absolute numbers of LKS (**E**) and LK (**F**) cells in control and infected mice at day 8, 11, 15, 24, 29 and 60 p.i.. **G**. Normalized percentages of LKS and LK cells within the Lin^-^ Kit^+^ population in control and infected mice at day 8, 11, 15, 24, 29 and 60 p.i.. In **B, C** and **E-G** data are presented as mean +/-s.e.m., and p values (asterisks) were determined by unpaired two-tailed Student’s *t*-test. * p < 0.05, ** p < 0.01, *** p < 0.001. *n* = 6, 9, 9, 9, 6, 6 control and *n* = 5, 7, 7, 8, 6, 6 infected mice at day 8, 11, 15, 24, 29, 60 p.i. respectively, pooled from three independent infections. PB, peripheral blood; BM, bone marrow; RBC, red blood cells; WBC, white blood cells, LK, Lineage^-^Ckit^+^Sca-1^-^, LKS, Lineage^-^Ckit^+^Sca-1^+^.

Acute infection was accompanied by anemia as shown by a significant decrease in the count of PB red blood cells (RBCs) at day 11, which gradually recovered and returned to normal levels by day 29 p.i. (Figure 1C, left panel). We observed different dynamics for white blood cells (WBCs): while they significantly decreased at day 11 p.i., they recovered faster than RBCs and overall presented larger variability within the cohort. Interestingly, at day 15 p.i., WBCs showed a trend to increase, which would be consistent with the activation of the immune system elicited by *P. chabaudi* ^19^ (Figure 1C, middle panel). Lastly, platelet counts were largely stable throughout the infection, with only a small but significant decrease in platelet numbers in infected animals at day 15 p.i. (Figure 1C, right panel).

To validate that the acute phase of *P. chabaudi* infection is associated with perturbed hematopoiesis, we analyzed BM composition using flow cytometry. First we observed broad changes in the overall primitive HSPC and oligopotent progenitor compartments, where the first compartment includes HSCs and MPP populations and the second one includes common myeloid progenitor and bipotent myeloid progenitors (Figure S1 and Figure 1D). At the peak of infection (day 11 p.i.), we observed an expansion of the LKS (Lineage^-^c-kit^+^Sca-1^+^ cells) compartment, which comprises MPPs and HSCs, in agreement with previous studies ^31^ (Figure 1D-E). This infection-induced increase in LKS cells absolute number was still observable at day 15 p.i., albeit at lower levels, and LKS cell counts had returned to control levels during early recovery (day 24 p.i.). The changes in the LKS population were mirrored by a trend for contraction and recovery of the more differentiated, myeloid-primed LK (Lineage^-^ c-kit^+^ Sca-1^-^) compartment, which was not statistically significant due to higher variability in the size of this population (Figure 1F). However, when the whole Lineage^-^ c-kit^+^ compartment (LK and LKS cells combined) was considered, the proportion of LKS cells within the compartment was strikingly increased, despite it normally being the smaller component (Figure 1D and G). For both LKS and LK compartments, counts recovered by day 24 and remained essentially unchanged from then (Figure S2A). Taken together, these data show that *P. chabaudi* is an effective model to study changes in HSPC populations in response to an acute infection and during recovery from it.

### Infection-induced alterations in the size of primitive hematopoietic stem and progenitor populations are resolved upon pathogen clearance

To investigate *P. chabaudi* infection-induced changes in primitive hematopoietic stem and progenitor cells, we examined subpopulations within the LKS compartment by employing signaling lymphocyte activation molecule (SLAM: CD150 and CD48) and FLK2 (CD135/Flt3) markers. In this study, we define MPP4 as LKS Flk2^+^ CD150^-^ cells, and MPP3 and MPP2 as LKS Flk2^-^ CD150^-^ CD48^+^ and LKS Flk2^-^ CD150^+^ CD48^+^ respectively, while MPP5-6 are defined as LKS Flk2^-^ CD150^-^ CD48^-/low^, according to previous studies ^3,32,33^. Lastly, we analyzed separately the more quiescent and primitive CD48^-^ HSC population, defined as LKS Flk2^-^ CD150^+^ CD48^-^, and the more proliferative, less primitive CD48^low/-^ HSPCs (LKS Flk2^-^CD150^+^ CD48^low/-^), as we described previously ^34^ (Figure 2, Figure S2). Consistent with the hallmarks of multiple inflammatory models^5,7,12,25^, *P. chabaudi* caused a dramatic swelling of all MPP populations, which contributed to the overall LKS population expansion observed earlier, while the most primitive CD48^-^ HSCs appeared reduced (Figure 2A).Overall, populations returned to levels similar to controls by day 24 p.i., followed by a small enlargement of MPP populations at day 29 p.i., which was then recovered at day 60 p.i. (Figure 2A, Figure S2B-C).

**Figure 2.**
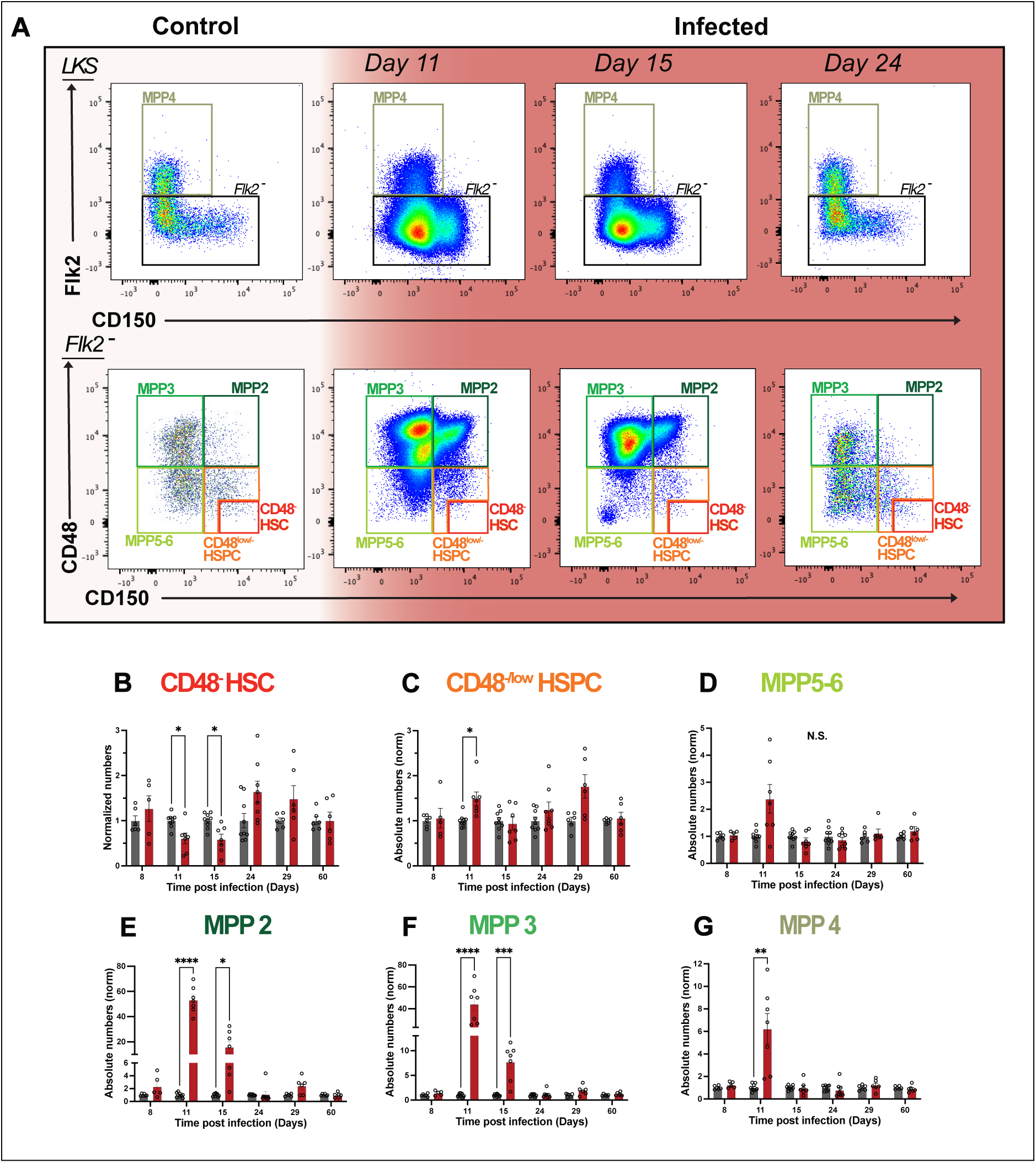
*P. chabaudi* infection effects on primitive hematopoietic populations are resolved upon pathogen clearance. **A.** Representative flow cytometry plots showing primitive HSPC populations in control and infected mice at day 11, 15 and 60 p.i.. **B-G.** Normalized absolute numbers of CD48^-^ HSC (**B**), CD48^low/-^ HSPC (**C**), MPP5-6 (**D**), MPP2 (**E**), MPP3 (**F**) and MPP4 (**G**) cell populations in control and infected mice at day 8, 11, 15, 24, 29 and 60 p.i.. In **B-G** data are presented as mean +/-s.e.m., and p values (asterisks) were determined by unpaired two-tailed Student’s *t*-test. * p < 0.05, ** p < 0.01, *** p < 0.001. Data were pooled from three independent infections. *n* = 6, 9, 9, 9, 6, 6 control and *n* = 5, 7, 7, 8, 6, 6 infected mice at day 8, 11, 15, 24, 29, 60 p.i. respectively, pooled from three independent infections. HSC, hematopoietic stem cell, HSPC, hematopoietic stem and progenitor cells, MPP, multipotent progenitor.

To quantify the precise effect of *P. chabaudi* infection on all the above-mentioned populations, we calculated and analyzed their absolute numbers (Figure 2B-G). The absolute number of CD48^-^ HSCs halved compared to controls during the acute phase of infection (days 11 and 15 p.i.), and showed a highly variable rebound phase, with a trend to increase at day 24 p.i., albeit not statistically significant. By day 60 p.i. CD48^-^ HSC numbers had returned to control levels (Figure 2B). Instead, CD48^low/-^ HSPC numbers were moderately but significantly increased at day 11 p.i., and equivalent to control levels on all other days, except for a trend towards an increase on day 29 p.i. (Figure 2C). The myeloid/lymphoid-primed MPP5-6 cells were quite stable, with only a non-significant trend to increase at day 11 p.i. (Figure 2D). As expected, the myeloid-primed MPPs populations, namely MPP2 and MPP3, had a massive increase at the peak of infection and subsequently recovered. Specifically, they were respectively 50 and 40 times more numerous compared to the control group at day 11 p.i. and returned to normal levels by day 24 (Figure 2E-F). Lymphoid-primed MPP4 cells showed a smaller, 6-fold increase in population size at day 11 p.i., which was followed by a quicker recovery, with baseline numbers reached by day 15 p.i. (Figure 2G). These data highlight that *P. chabaudi* infection-induced emergency granulopoiesis is fueled by a dramatic increase of myeloid-primed MPPs and by a decrease in quiescent HSCs. Although the size of these populations recovers upon pathogen disappearance, the question remains on the proliferation dynamics that underpin the changes observed during the acute and recovery phases of infection.

### Heterogeneous proliferation dynamics within the primitive haematopoietic compartment during and following infection

To shed light on possible mechanisms underlying the increase and decrease of primitive hematopoietic cell numbers, we assessed the proliferation of these populations by measuring *in vivo* EdU incorporation. EdU is a thymidine analogue that intercalates in the DNA of cells during S-phase of the cell cycle and can be detected *ex vivo* using flow cytometry. To gain a quantitative and comprehensive understating of proliferation changes in HSPCs we assessed both absolute number and percentage of proliferating (EdU^+^) cells within each primitive population (Figure 3, Figure S3). While the absolute numbers are useful to have an overall quantification, percentages are key to understand whether the increase or decrease in the absolute numbers of cell populations results from an increase or decrease in their proliferation rate. Because CD48^-^ HSCs are very rare and quiescent, flow cytometry data were not sufficiently robust to quantify the absolute number of proliferating cells. However, we consistently detected more numerous EdU^+^ cells in this population at day 11 and 15, but not at any other time point analyzed (Figure 3A, Figure S3A). Consistent with this, we detected a significant increase in the percentage of EdU^+^ CD48^-^ HSCs during the acute phase of infection, which returned to control levels by day 24 p.i. (Figure 3B). Similarly, both absolute number and percentage of proliferating cells within CD48^low/-^ HSPCs were increased at infection peak and returned to control levels by day 15 p.i and day 24 p.i., respectively (Figure 3C).

**Figure 3.**
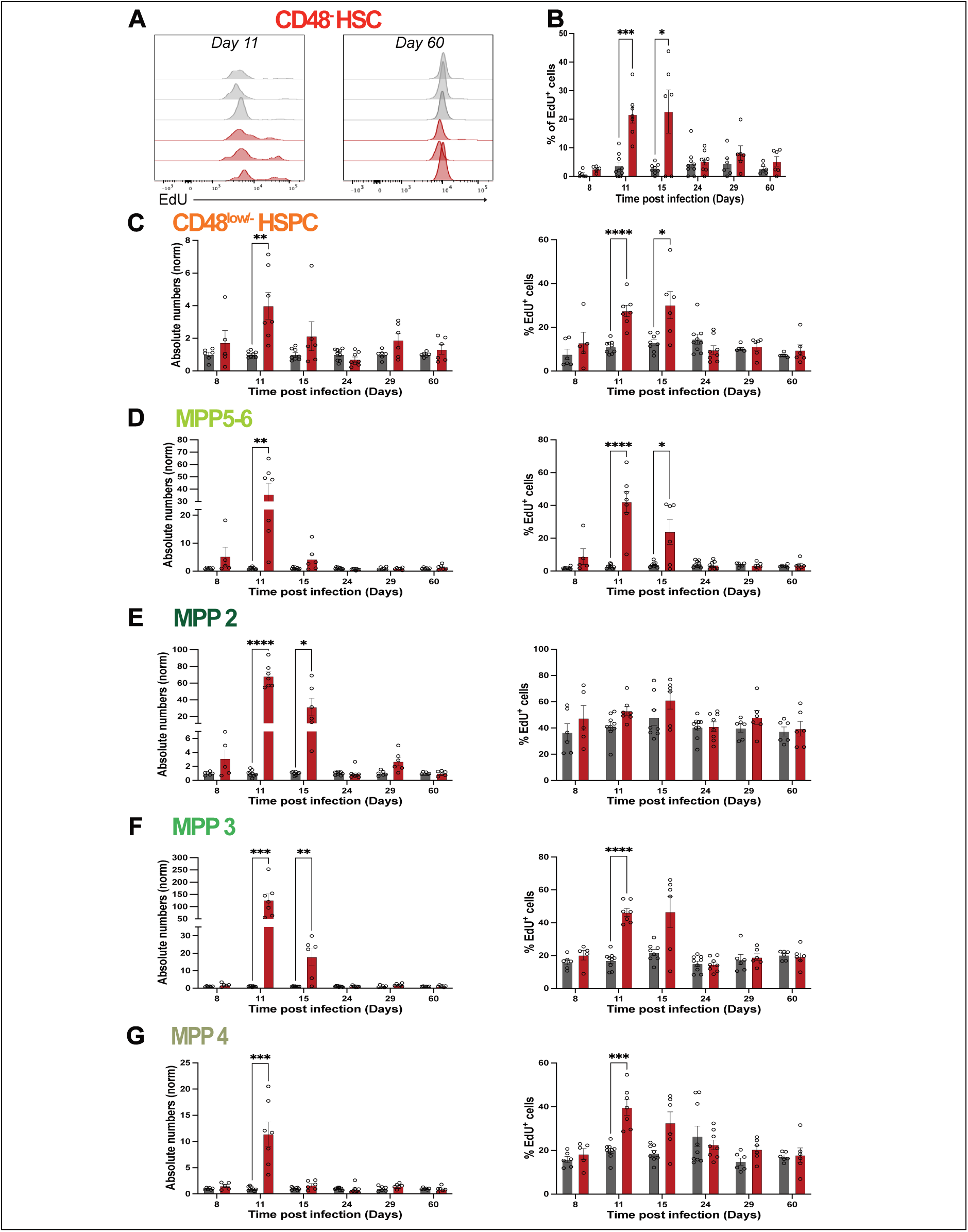
Proliferation dynamics of primitive HSPC populations during *P. chabaudi* infection. **A.** Representative flow cytometry data histograms showing EdU incorporation in CD48^-^ HSCs in control and infected mice at day 11 and 60 p.i. (each histogram represents one mouse, plots are representative of all time points analyzed from 3 independent infections). **B.** Percentage of EdU^+^ cells in CD48^-^ HSCs in control and infected mice at day 8, 11, 15, 24, 29 and 60 p.i.. **C-G.** Normalized absolute numbers (left) and percentages (right) of EdU^+^ cells in CD48^low/-^ HSPC (**C**), MPP5-6 (**D**), MPP2 (**E**), MPP3 (**F**), and MPP4 (**G**) populations in control and infected mice at day 8, 11, 15, 24, 29 and 60 p.i.. In **B-G** data are presented as mean +/-s.e.m., and p values (asterisks) were determined by unpaired two-tailed Student’s *t*-test. * p < 0.05, ** p < 0.01, *** p < 0.001. Data were pooled from three independent infections. *n* = 6, 9, 9, 9, 6, 6 control and *n* = 5, 7, 7, 8, 6, 6 infected mice at day 8, 11, 15, 24, 29, 60 p.i. respectively.

MPP populations presented more heterogeneous kinetics (Figure 3D-G). Interestingly, while the absolute numbers of MPP5-6 only showed a trend to increase (Figure 2D), the number of EdU^+^ cells in this population was 35 times higher in infected animals compared to control animals at day 11 p.i. (Figure 3D, left panel, Figure S3C). Similarly, percentages of EdU^+^ cells within MPP5-6 increased from 2.8% in control to 42% at day 11 p.i. (Figure 3D, right panel). However, while numbers recovered quickly, already by day 15 p.i., percentages of proliferating cells within these populations returned to control levels only at day 24 p.i. (Figure 3D, right panel). When analyzing MPP2, 3 and 4 populations, we noticed that the expansion of cell numbers observed during the acute phase of infection (Figure 2E-G) was associated with dramatic increase in the absolute number of EdU^+^ cells within all the populations, but with cell type specific dynamics in the proportion of EdU^+^ cells and in the time of recovery (Figure 3E-G, Figure S3C-F). In detail, the massive expansion observed in megakaryocytic/erythroid-biased MPP2 at peak of infection (Figure 2E) was accompanied by a 68-fold increase in the numbers of proliferating cells within this compartment, which recovered by day 24 p.i. (Figure 3E, left panel). Interestingly, no changes were observed in terms of percentages of proliferating cells within MPP2 throughout the infection (Figure 3E, right panel). Similarly, at day 11 p.i., the enlargement of the myeloid-biased MPP3 compartment was also accompanied by a massive increase in the absolute numbers of proliferating cells within this population compared to controls (120 times higher), which then returned to homeostatic levels by day 24 p.i. (Figure 3F, left panel). MPP3s’ proportion of EdU^+^ cells also increased, though only about 2.5 fold, and recovered already by day 15 p.i. (Figure 3F, right panel). Lastly, while lymphoid-biased MPP4s showing a 6-fold population enlargement at day 11 p.i. (Figure 2E), absolute numbers of proliferating cells within this compartment were 12-fold higher compared to controls at day 11 p.i., and returned to control levels by day 15 p.i. (Figure 3G, left panel). Similarly, percentages of proliferating cells within MPP4s also peaked at day 11 p.i. and were back to homeostasis at day 15 p.i. (Figure 3G, right panel). Taken together, these data demonstrate that *P. chabaudi* infection leads to a significant but transient increase in proliferating cells within the primitive hematopoietic compartments, including CD48^-^ HSCs, which is then followed by recovery dynamics specific for each population.

### *P. chabaudi* infection dramatically affects oligopotent progenitor numbers

To gain a fuller picture of the changes occurring to different hematopoietic progenitor populations in response to malaria infection we extended our flow cytometry analysis and investigated the numbers and proliferative state of MPPs’ immediate progeny. These populations are known to be lineage restricted, and therefore oligopotent. and more proliferative than primitive HSPCs^35^. Guided by well-established definitions based on the expression of CD34 and CD16/32 receptors, we defined CMPs, GMPs and MEPs within the LK compartment as CD34^+^ CD16/32^-^, CD34^+^ CD16/32^+^ and CD34^-^ CD16/32^-^ respectively (Figure 4A, Figure S4A)^36^. Lastly, CLPs were defined as Lin^-^ c-kit^int^ Flk2^+^ CD127^+^ (Figure 4B, Figure S4B)^7^. We observed a drastic decrease in the CMP, GMP, MEP and CLP populations at the peak of the infection, and again population-specific patterns of recovery emerged at the later time points (Figure 4). Specifically, CMPs’ absolute numbers decreased to nearly zero at day 11 p.i, were still reduced compared to controls at day 15 and returned to homeostatic levels only by day 29 p.i. (Figure 4C, Figure S4A). GMP and MEP numbers in infected mice were reduced by 75% at day 11 p.i. compared to control mice, however recovery was already observed at day 15 p.i. for MEPs, while it was reached by day 24 p.i. for GMP (Figure 4D, E). Interestingly, despite CLP numbers being reduced to nearly zero at day 11 p.i., control levels were quickly re-established by day 15 p.i. (Figure 4F). Additionally, we included the LK CD34^-^ CD16/32^+^ population in our analysis, which is negligible in steady state, but we noticed was increased during infection (Figure 4G). This population has been generally neglected and is therefore poorly caracterized. The size of this population increased about 2-fold only by day 15 p.i., and returned to steady state size by day 24 p.i. (Figure 4G). Overall, these data demonstrate that despite a general loss of oligopotent progenitors within the BM compartment during *P. chabaudi* infection, their numbers are recovered within one month p.i., with the CMP population being the slowest to recover and thus being the most affected.

**Figure 4.**
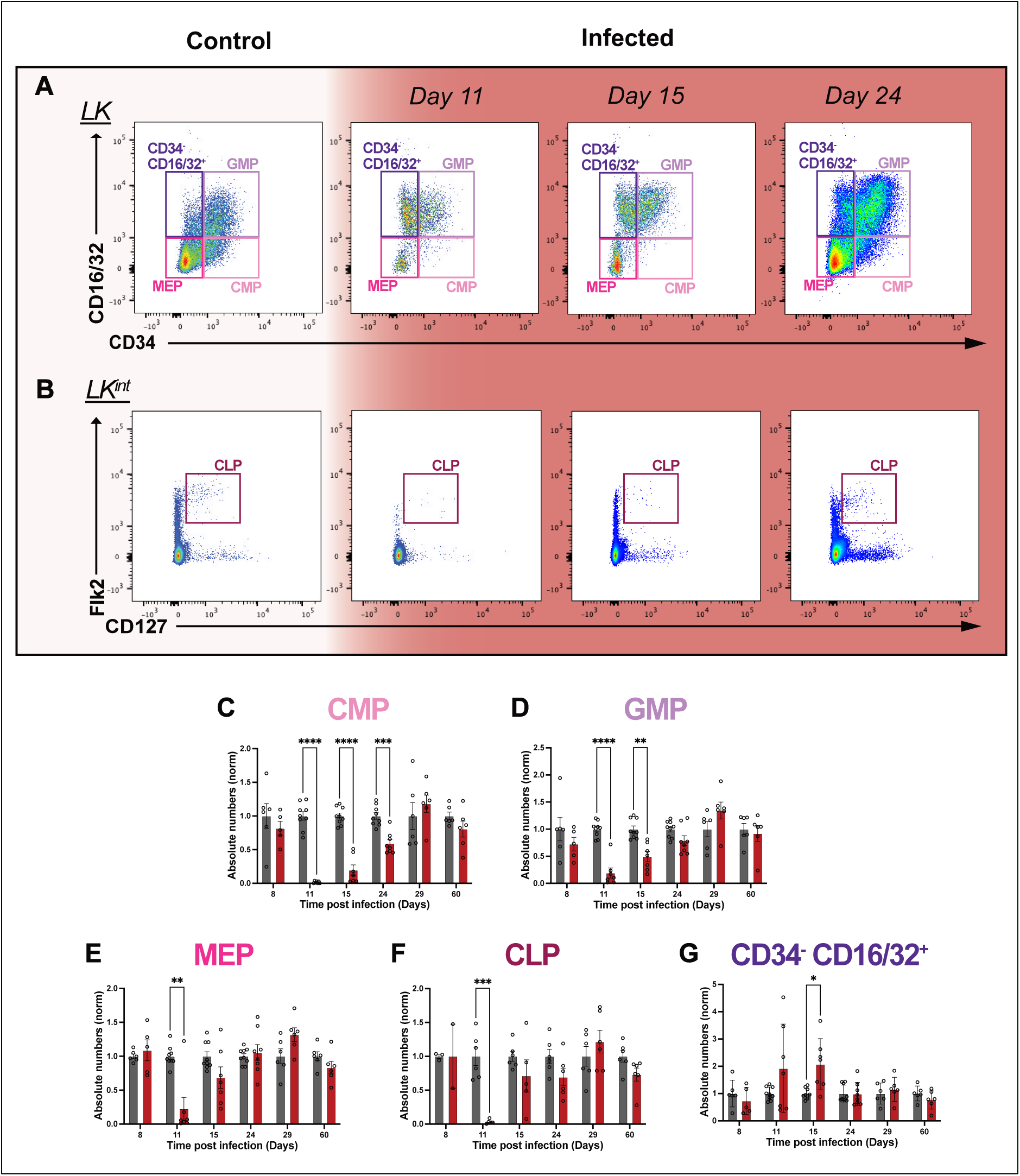
Mature progenitors are largely lost during acute *P. chabaudi* infection. **A-B**. Representative flow cytometry plots showing CD34^-^CD16/32^+^, GMP, CMP and MEP LK subpopulations (**A**) and CLPs (**B**) in control and infected mice at day 11, 15 and 60 p.i. **C-G.** Normalized absolute numbers of CMP (**C**), GMP (**D**), MEP (**E**), CLP (**F**), and CD34^-^CD16/32^+^ (**G**) progenitors in control and infected mice at day 8, 11, 15, 24, 29 and 60 p.i. In **C-G** data are presented as mean +/-s.e.m., and p values (asterisks) were determined by unpaired two-tailed Student’s *t*-test. * p < 0.05, ** p < 0.01, *** p < 0.001. *n* = 6, 9, 9, 9, 6, 6 control and *n* = 5, 7, 7, 8, 6, 6 infected mice at day 8, 11, 15, 24, 29, 60 p.i. respectively. LK^int^, Lineage^-^ c-Kit^intermediate^; CMP, common myeloid progenitor; GMP, granulocyte-macrophage progenitor; MEP; megakaryocyte-erythrocyte progenitor; CLP, common lymphoid progenitor.

### Oligopotent progenitors’ proliferation is minimally affected during *P. chabaudi* infection

To explore how mature progenitors’ numbers respond to and recover following *P. chabaudi* infection we investigated their proliferation levels throughout infection and recovery. Similar to our analysis of primitive HSPCs, we assessed both absolute number and percentage of proliferating cells within each population, building a comprehensive overview of the dynamics taking place at this stage of hematopoietic development (Figure 5, Figure S5). Predictably, the overall decrease in the absolute numbers of mature progenitors was accompanied by an overall decrease in the absolute numbers of proliferating cells within most subpopulations (Figure 5 A-E, left panels). Surprisingly, the reduction in proliferating cells’ numbers was not associated with either a reduction or an increase in the proportion of proliferative cells in any of the populations, as for the most part we did not observe any significant change in the percentages of proliferating cells within the subpopulations (Figure 5 A-E, right panels). As expected, in homeostasis (control mice), we found that most mature progenitors, namely GMPs, MEPs, and CD34^-^ CD16/32^+^ cells were associated with a very high proliferative index as approximately 40-50% of cells within each population were in S-phase (Figure 5 B, C and E, right panels). Interestingly, CMPs and CLPs were found to have a lower, but still relatively high proliferative index, with an average of 20% of proliferating cells within each population (Figure 5 A and D, right panels).

**Figure 5.**
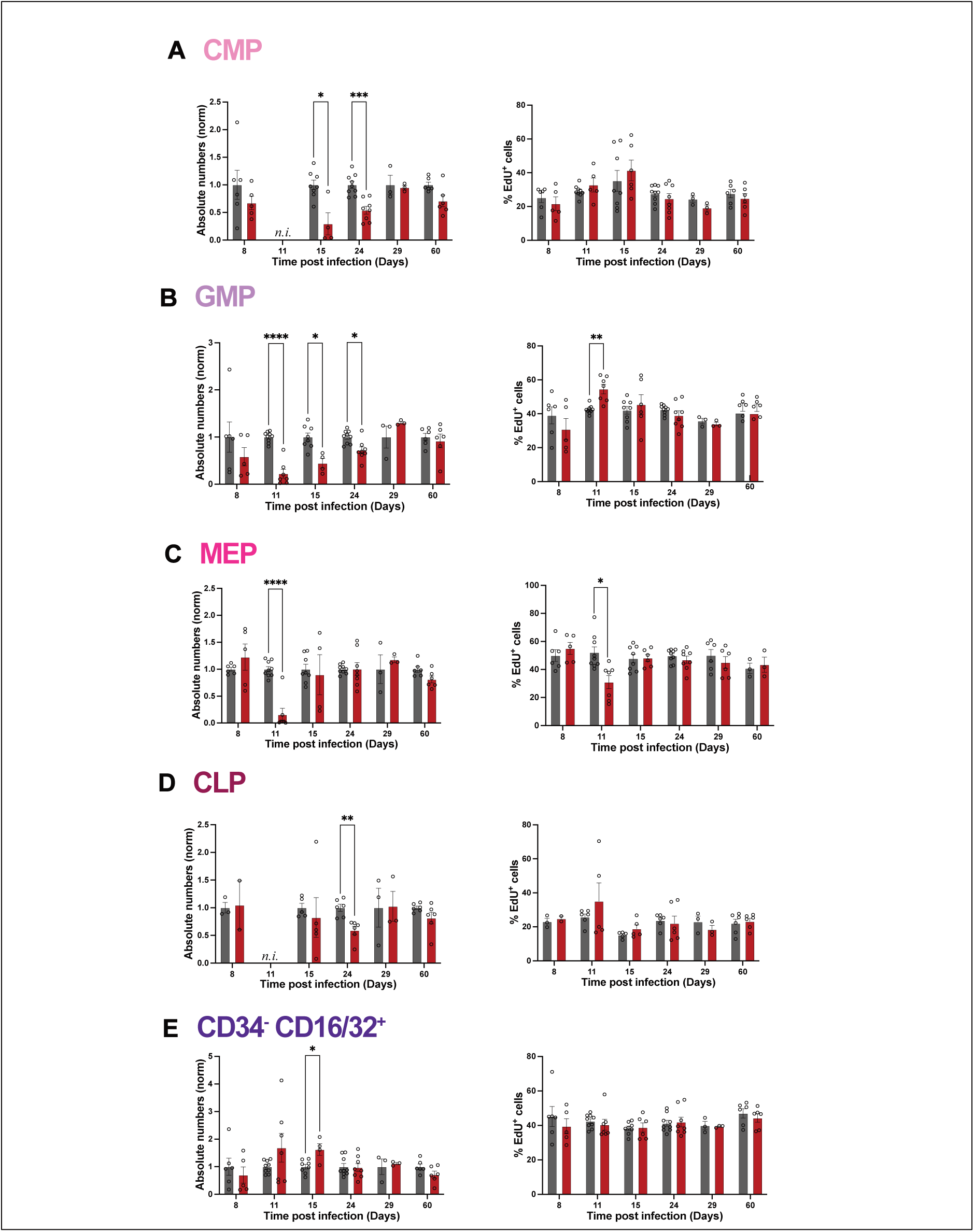
*P. chabaudi* infection does not alter the proliferation of mature progenitors. **A-E.** Normalized absolute numbers (left) and percentages (right) of EdU^+^ cells in CMP (**A**), GMP (**B**), MEP (**C**), CLP (**D**) and CD34^-^ CD16/32^+^ (**E**) populations in control and infected mice at day 8, 11, 15, 24, 29 and 60 p.i.. All data are presented as mean +/-s.e.m., and p values (asterisks) were determined by unpaired two-tailed Student’s *t*-test. * p < 0.05, ** p < 0.01, *** p < 0.001. Data were pooled from three independent infections experiments. *n* = 6, 9, 9, 9, 6, 6 for control and *n* = 5, 7, 7, 8, 6, 6 for infected mice at day 8, 11, 15, 24, 29, 60 p.i. respectively. n.i., not included.

A detailed analysis of the dynamics of each progenitor population uncovered again specific responses. Due to the virtual disappearance of the CMP population at day 11 p.i., their proliferation could not be assessed on that day. However, absolute numbers of proliferating cells within CMP were decreased on days 15 and 24 p.i., recovering by day 29 p.i. (Figure 5A, left panel, Figure S5A). Predictably, this recovery was in line with overall absolute numbers of CMPs reaching control levels by day 29 p.i. (Figure 4C). Instead, percentages of proliferating cells within this population remained similar to those of control mice throughout the infection (Figure 5A, right panel). Similarly to CMPs, numbers of proliferating GMPs also decreased at acute infection, recovering at day 29 p.i. (Figure 5B, left panel, Figure S5B). Of note, percentages of proliferating GMPs increased at day 11 p.i., but were similar to those of GMPs from control animals for all other time points analyzed (Figure 5B, right panel). Next, absolute numbers of proliferating MEPs decreased at day 11 p.i. (Figure 5C, left panel), coupled to a decrease in the proportion of proliferating cells within the remaining population (Figure 5C, right panel). MEP numbers and proportions of proliferative cells were similar to controls for all other time points (Figure 5C). Interestingly, with regards to CLPs, the absolute number of proliferating cells within this population was lower than control only at day 24 p.i. (day 11 data was not included as CLP were virtually absent at this timepoint) (Figure 5D, left panel). Similar to the other populations, the percentage of proliferating cells did not show any alteration (Figure 5D, right panel). Strikingly, when investigating the CD34^-^ CD16/32^+^ population we observed a very different response as there was a trend to increase in the number of proliferating cells at the peak of parasitaemia, day 11 p.i., with high mouse-to-mouse variability, followed by a significant increase compared to control at first time point post parasitemia peak, day 15 p.i., and a quick recovery, already achieved by day 24 p.i. (Figure 5E, left panel). However, no changes were detected in the percentage of proliferating cells throughout our analysis (Figure 5E, right panel). Taken together, these data demonstrate that committed hematopoietic progenitor populations maintain a relatively steady proliferative rate in response to *P. chabaudi* infection and subsequent recovery.

### Primitive and committed hematopoietic progenitor populations respond to and recover from *P. chabaudi* infection differently

To gain a system-level understanding of the cellular dynamics driving the hematopoietic response to and recovery from *P. chabaudi* infection, we combined all our observations in summary graphs where we could more effectively compare population dynamics (Figure 6). This analysis highlighted stark differences in the response of different cell populations. Within the primitive hematopoietic compartment, CD48^-^ HSCs were the only population with a significant reduction in size at day 11 and 15 p.i., and their downstream progenitor populations showed from minimal to very dramatic increase in size at day 11 p.i., with MPP 2 and 3 taking longer to return to control size (Figure 6A). Despite no statistically significant differences being highlighted for any of these populations after day 15 p.i., it was striking that all populations were synchronized in showing a trend to increase at day 29 p.i., likely due to the small increase in parasitemia, and again steady-state sizes at day 60 p.i. (Figure 6A). The overview of proliferation dynamics highlighted how the increase in MPP’s numbers was fueled by increased proliferation in all populations, albeit again at different degrees, and with the most primitive population, CD48^-^ HSCs, showing one of the highest increases in proliferation, likely to fuel activation of the hematopoietic cascade and regeneration of downstream progenitors (Figure 6B). Interestingly, while percentages of proliferating cells for most MPPs peaked with parasitemia at day 11 p.i., CD48^-^ HSCs had the highest proportion of EdU^+^ cells at day 15 p.i. (Figure 6B).

**Figure 6.**
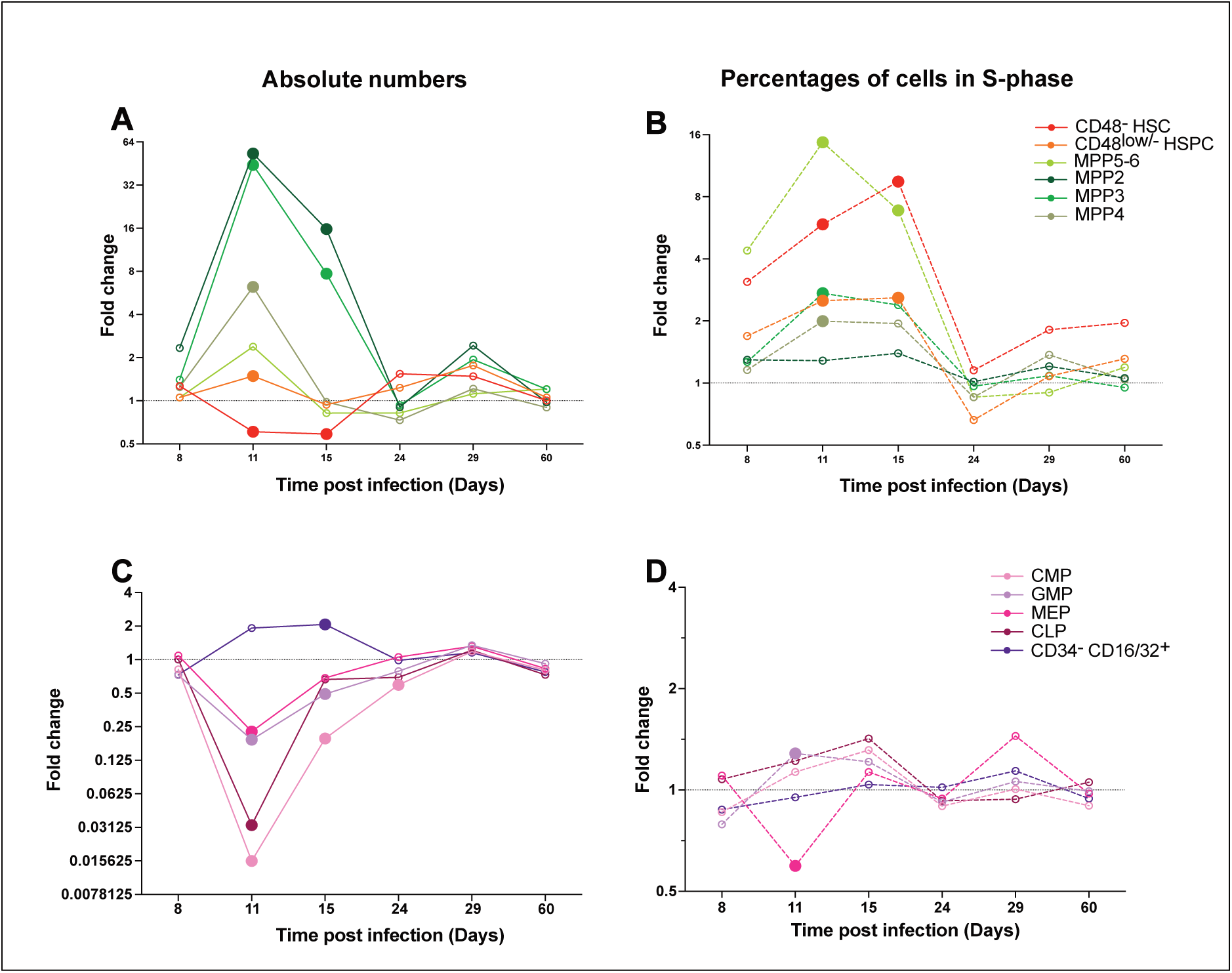
Hematopoietic populations adopt very different responses when responding to and recovering from severe infection. **A.** Average fold change of normalized absolute numbers of primitive hematopoietic cell populations. **B.** Average fold change of percentages of EdU^+^ cells in primitive HSPC populations. **C.** Average fold change of normalized absolute numbers of oligopotent progenitor populations. **D.** Average fold change of percentages of EdU^+^ cells in oligopotent progenitor populations. Data shown are a transformation of data showed in figures 2-5. 1 = average control values; values between 0 and 1 represent a decrease; value > 1 represent an increase. Statistically significant differences between infected and control group are indicated by bigger, filled circles. Non statistically significant differences are indicated by smaller, empty circles. Data were pooled from three independent infections experiments. *n* = 6, 9, 9, 9, 6, 6 control and *n* = 5, 7, 7, 8, 6, 6 infected mice at day 8, 11, 15, 24, 29, 60 p.i. respectively.

On the other hand, downstream oligopotent, committed progenitors presented a very different picture, with most populations massively reducing in size at the peak of parasitemia (day 11 p.i.), starting to recover steadily by day 15 p.i., and showing virtually no rebound in numbers at later time points (Figure 6C). Grouped analysis confirmed the overall lack of dramatic changes in proliferation for all these populations, the only exception being MEPs, which showed a trend to reduced proliferation at day 11 p.i. (Figure 6D). All these data together demonstrated how different populations within the hematopoietic tree adopt very different dynamics when responding to and recovering from infection. Generally, while primitive progenitors massively increased their population size as well proliferation levels, mature progenitor populations were largely affected through a high decrease in size but no significant changes in proliferation rate. Moreover, quiescent HSCs were shown to respond to infection by decreasing in number at peak infection followed by an increase in percentages of proliferating cells. For the most part, recovery, in terms of population size and proliferation, was observed within one month p.i..

## DISCUSSION

Many studies have described the effects of acute infection on the primitive compartment of the hematopoietic system, including an increase in proliferation, increase in myeloid differentiation and an increase in mobilization. However, numerous questions regarding the recovery of the hematopoietic system following inflammatory stimuli are still unanswered, such as when, how and to what extent the system recovers. Additionally, the effects of infection on oligopotent progenitor cell populations have remained overlooked. For this study, we employed a non-lethal model of malaria*, P. chabaudi*, from which mice can recover without pharmacological interventions, thus allowing us to characterize the dynamics of different hematopoietic populations during the acute and recovery phases of a blood-stage infection, without any potential combinatorial effects by drug treatments. By systematically tracking the changes in the size of all primitive hematopoietic compartments we could identify unique patterns of hematopoietic remodeling and return to homeostatic levels.

A hallmark of many bacterial and viral infections, as well as infections caused by *Plasmodium* species, is increased proliferation within the primitive hematopoietic compartments^6,37,38^, and systematic analysis of the proliferation of primitive hematopoietic populations responding to and recovering from *P. chabaudi* infection provided mechanistic insights into populations’ increase and decrease and revealed again unique dynamics. Measuring EdU uptake after a single pulse of drug administration provides a qualitative measurement of changes in proliferation rates of each cell population of interest, and a double pulse of two different thymidine analogues is required to calculate exact proliferation rates^34^. Nevertheless, comparison of the frequency of EdU^+^ cells per population between control and infected mice and between different populations provides a simple mean of uncovering relative changes in the proliferation rate of HSPC populations. Namely, the most primitive CD48^-^ HSCs were detected to decrease in numbers but increase proliferation at the peak of infection, consistent with reports studying other infection models^16,25,39,40^. MPP populations dramatically increased in size and proliferation, while oligopotent progenitors virtually disappeared at the peak of the infection but their proliferation was never altered. This indicates that primitive hematopoietic populations adopt very different dynamics in response to a natural infection. Lastly, we demonstrated that recovery of population size and proliferation within the hematopoietic system occurred within one month p.i.. Interestingly, mouse to mouse variability made it challenging to ascertain whether any rebound, which would most often be expected when any perturbed system returns to steady state, would consistently take place. The ability to respond rapidly to *Plasmodium* infection and to return to homeostatic hematopoiesis as the infection is resolved is a clear indication of the ability of the organism to cope with such infection, which mice and humans have evolved with. Moreover, our datasets can be interrogated to start addressing whether there may be any hematopoietic long-term consequences of surviving *Plasmodium* infection.

It has been widely recognized that replacement of consumed immune cells during an infection depends on hematopoiesis, which is remodeled by the ongoing immune responses in order to replenish lineage-specific cells which are key in clearing a particular pathogen. Plasticity in the size of primitive hematopoietic populations is key in supporting hematopoiesis ‘on demand’. For instance, malaria has been widely associated with a switch to myelopoiesis at the expense of lymphopoiesis as innate immune cells are key in the efficient destruction of parasitized RBCs ^31,41^. This is further supported by our data as we observed a deeper effect on the CMP population, which showed both the most dramatic decrease and slowest recovery compared to the other oligopotent precursors. Both the dramatic decrease and slow recovery of population size may be the result of the rapid induction and likely persistence of fast differentiation of any cells reaching the CMP compartment. The rapid recovery of MEP and CLP numbers likely reflects the evolutionary requirement to restore healthy hematopoiesis as quickly as possible following infection, however, the different rates of recovery of different oligopotent progenitor populations may be a possible cause of some immunological abnormalities detected for a number of weeks in malaria survivors^42,43^.

Interestingly, we did not detect a change in the proliferation rate of all oligopotent progenitors, apart from a moderate increase in proliferation of GMPs at day 11 p.i.. In homeostasis oligopotent progenitors had the highest proliferative rate, which is consistent with loss of self-renewal and increased lineage commitment being associated with increased proliferation, leading to downstream ‘bursts’ of differentiated cells production^44–46^. The overall lack of changes in proliferation rate of oligopotent progenitors is not limited to *Plasmodium* infection. Walters et al.^47^ previously demonstrated that oligopotent progenitors were unresponsive to polyinosinic:polycytidylic acid (pI:pC) and that they were associated with a higher proliferative index during homeostasis compared to LT-HSCs. Oligopotent progenitors’ proliferation rates might be fixed at the homeostatic levels for yet unexplained reasons. These cells might already proliferate at their maximum capacity during homeostasis, which is supported by our study as we reported percentages of proliferating cells within mature progenitors to reach 40% and even 60%, while in the primitive compartments, they were below 20%. Alternatively, *Plasmodium* induced stress might not be sufficient to elicit proliferation of these cells, which might have a higher threshold of activation, or only respond to different types of stress altogether.

The MPP compartment was undoubtedly the one most remodeled by *Plasmodium* infection, with dramatic changes in both population sizes and proliferative state. In homeostasis, MPP4 cells with both lymphoid and myeloid potential ensure balanced blood cell output ^32,48^. Simultaneously, lower numbers of MPP2 and MPP3 cells, associated with megakaryocyte/erythroid and myeloid output respectively, contribute to daily hematopoiesis^32^. Interestingly, in steady state, MPP2 cells displayed the highest percentages of proliferating cells (40%) compared to all the other MPP populations (<20%), consistent with the rapid reconstitution ability that has been previously associated with MPP2 cells ^48^. Upon inflammatory challenges that trigger emergency myelopoiesis, instructive differentiation signals are activated within the primitive compartment and lead to the expansion and skewing of the MPP compartment^32,49,50^. Consistent with this literature, our data showed a massive expansion of MPP2 and MPP3 populations, which was then recovered within day 24 p.i.. This expansion was supported by a dramatic increase in the proliferation rate of most MPP populations, especially the MPP5/6 population, which is hierarchically located between HSCs and MPP2-4 ^33^. Despite the dramatic boost in proliferation, the numbers of MPP5/6 remained unchanged, underscoring the importance of this population in the production of the downstream MPPs, with cells transitioning very quickly through this compartment. On the contrary, the proliferation rate of megakaryocytic/erythroid-biased MPP2s remained relatively stable throughout our analysis, and at similar levels of oligopotent progenitors’ proliferation. Of note, it has been reported that following acute physiologic insult the skewing of hematopoiesis into the production of myeloid cells leads to the impairment of erythro-/megakaryopoiesis^51^. Specifically, inflammation-induced release of IL-1β or IFNγ has been shown to directly increase the expression of *Pu.1* on MPPs, which in turn stimulates myeloid production while inhibiting erythroid development ^17,52^. It is possible that in infected mice MPP2 cells were not stimulated to proliferate further, and that the enlargement of the population may result from impaired differentiation into downstream populations. Additionally, it is likely that MPP2 cells would be transcriptionally rewired to switch to a myeloid output, as it was previously described for IL-7Rα^+^c-Kit^high^ lymphoid progenitors^24^. Similarly, it is expected that the transcriptional rewiring of lymphoid committed progenitors likely already initiates in the MPP4 population.

Importantly, all primitive populations, including CD48^low^ HSPCs, MPP5/6 and MPP2, 3 and 4 returned to homeostatic size and proliferative rates by day 15 or 24 p.i., highlighting the great flexibility in their dynamics and output, and consistent with primitive progenitor populations responding directly to inflammatory stimuli rather than purely responding to a domino effect from consumption of differentiated cells^9^. The fact that anemia lasts longer than the disruption of HSPC populations is likely the compound result of RBC destruction by parasites and of the time required to rewire hematopoiesis to support erythropoiesis again, the latter driven by both transcriptional changes and microenvironmental stimuli.

Analysis of the most primitive hematopoietic stem cells, CD48-HSCs revealed an initial depletion of the population, which had been previously demonstrated in response to *P. berghei*^25^. Many factors could explain this consumption, and it is generally accepted that inflammation causes premature differentiation of HSCs at the expense of self-renewal^10,17^. In addition, apoptosis and mobilization outside of the BM constitute other possible mechanisms leading to HSC disappearance. We demonstrated that the proportion of proliferating CD48^-^ HSCs increased during the acute stages of infection, consistent with previous work with *P. berghei* ^7^ and other models of inflammatory stress^6,12,53,54^, and feeding the increase of the MPP compartment. Interestingly, CD48-HSC numbers were on average back to homeostatic levels by day 24 p.i., however remained highly variable compared to those of all other primitive populations and may reflect mouse-to-mouse variability in the return to homeostasis for the most primitive HSCs. Many mechanisms are in place to control the proliferation and differentiation of HSPCs, including feedback signals from neighboring cells or from the specialized HSC niche ^55^. In addition, HSCs can sense the presence of infectious agents directly, and human *Plasmodium* species have been shown to take residence in the BM ^56^. It is possible that the observed variable numbers of CD48^-^ HSCs are a sentinel of a similar re-localization by *P. chabaudi.* Consistent with this, we observed a trend to a slight increase in numbers of all primitive HSPC populations at the day 29 p.i. time point, when a small recrudescence of *P. chabaudi* growth has been reported ^30^.

The recovery of HSC numbers following their decrease at the peak of infections highlights the robustness of the system, however, it raises the question whether the recovered HSC population may be less heterogeneous and/or less polyclonal than that of mice that remain healthy. This has important implications for the potential development of clonal hematopoiesis following severe or repeated infections. This would be highly relevant for the human population of endemic malaria regions, which could be at higher risk of developing hematological malignancies such as myeloid leukemia. It has already been shown that DNMT3a mutant HSCs have a selective advantage in surviving high levels of IFNγ^15^, a key upregulated cytokine in malaria infection^25,31,57,58^, but repeated, infection-induced bottlenecks in the HSC population may lead to a reduction in HSC clonality even independently of pre-leukemic mutations, and this would inherently affect the robustness of the overall hematopoietic system long-term.

In conclusion, our study indicates that HSCs are activated and depleted during *P. chabaudi* infection, which is consistent with the accepted paradigm of HSCs adjusting progeny output. Meanwhile, the oligopotent progenitors are quickly consumed to compensate the consumption of short-lived immune cells, and the main hematopoietic output burden relies on the proliferation and differentiation abilities of MPPs, which serve as the ideal valve for rapidly generating cells. Overall, despite the substantial perturbation we documented following *P. chabaudi* infection, hematopoietic homeostasis is recovered one month p.i.. All together these data demonstrate the high heterogeneity present within the hematopoietic system and outline the behaviors specific to each hematopoietic population in response to an infection. Importantly, this study identifies a window of time following *Plasmodium chabaudi* infection when recovery could be improved by boosting the hematopoietic restoration process. For instance, targeted stimulation of MPPs to produce erythroid precursors could rescue impaired erythropoiesis, leading to amelioration of severe anaemia. Additionally, enhancing mature progenitors’ proliferation could reduce their depletion. Crucially, accelerating hematopoietic recovery would improve the return of red and immune cells to homeostatic levels and could minimize the risk of co-infection, which is often observed in malaria patients^59^. Our findings highlight the significance of gaining a deeper understanding of the mechanisms involved in responding to and recovering from infection to prevent the long-term effects associated with this disease.

## REFERENCES

1. Caiado F, Pietras EM, Manz MG. Inflammation as a regulator of hematopoietic stem cell function in disease, aging, and clonal selection. J Exp Med. 2021;218(7):e20201541. doi:10.1084/jem.20201541

2. Sezaki M, Hayashi Y, Wang Y, Johansson A, Umemoto T, Takizawa H. Immuno-Modulation of Hematopoietic Stem and Progenitor Cells in Inflammation. Front Immunol. 2020;11:585367. doi:10.3389/fimmu.2020.585367

3. Purton LE. Adult murine hematopoietic stem cells and progenitors: an update on their identities, functions, and assays. Exp Hematol. 2022;116:1–14. doi:10.1016/j.exphem.2022.10.005

4. Kondo M, Wagers AJ, Manz MG, et al. Biology of Hematopoietic Stem Cells and Progenitors: Implications for Clinical Application. Annu Rev Immunol. 2003;21(1):759–806. doi:10.1146/annurev.immunol.21.120601.141007

5. Takizawa H, Fritsch K, Kovtonyuk LV, et al. Pathogen-Induced TLR4-TRIF Innate Immune Signaling in Hematopoietic Stem Cells Promotes Proliferation but Reduces Competitive Fitness. Cell Stem Cell. 2017;21(2):225–240.e5. doi:10.1016/j.stem.2017.06.013

6. Baldridge MT, King KY, Boles NC, Weksberg DC, Goodell MA. Quiescent hematopoietic stem cells are activated by IFNγ in response to chronic infection. Nature. 2010;465(7299):793-797. doi:10.1038/nature09135

7. Vainieri ML, Blagborough AM, MacLean AL, et al. Systematic tracking of altered haematopoiesis during sporozoite-mediated malaria development reveals multiple response points. Open Biol. 2016;6(6):160038. doi:10.1098/rsob.160038

8. Hormaechea-Agulla D, Le DT, King KY. Common Sources of Inflammation and Their Impact on Hematopoietic Stem Cell Biology. Curr Stem Cell Rep. 2020;6(3):96–107. doi:10.1007/s40778-020-00177-z

9. King KY, Goodell MA. Inflammatory modulation of HSCs: viewing the HSC as a foundation for the immune response. Nat Rev Immunol. 2011;11(10):685–692. doi:10.1038/nri3062

10. Bogeska R, Mikecin AM, Kaschutnig P, et al. Inflammatory exposure drives long-lived impairment of hematopoietic stem cell self-renewal activity and accelerated aging. Cell Stem Cell. 2022;29(8):1273–1284.e8. doi:10.1016/j.stem.2022.06.012

11. Pietras EM, Lakshminarasimhan R, Techner JM, et al. Re-entry into quiescence protects hematopoietic stem cells from the killing effect of chronic exposure to type I interferons. J Exp Med. 2014;211(2):245–262. doi:10.1084/jem.20131043

12. Essers MAG, Offner S, Blanco-Bose WE, et al. IFNα activates dormant haematopoietic stem cells in vivo. Nature. 2009;458(7240):904-908. doi:10.1038/nature07815

13. Wilson A, Laurenti E, Oser G, et al. Hematopoietic Stem Cells Reversibly Switch from Dormancy to Self-Renewal during Homeostasis and Repair. Cell. 2008;135(6):1118–1129. doi:10.1016/j.cell.2008.10.048

14. Esplin BL, Shimazu T, Welner RS, et al. Chronic Exposure to a TLR Ligand Injures Hematopoietic Stem Cells. J Immunol. 2011;186(9):5367–5375. doi:10.4049/jimmunol.1003438

15. Hormaechea-Agulla D, Matatall KA, Le DT, et al. Chronic infection drives Dnmt3a-loss-of-function clonal hematopoiesis via IFNγ signaling. Cell Stem Cell. 2021;28(8):1428–1442.e6. doi:10.1016/j.stem.2021.03.002

16. Matatall KA, Jeong M, Chen S, et al. Chronic Infection Depletes Hematopoietic Stem Cells through Stress-Induced Terminal Differentiation. Cell Rep. 2016;17(10):2584–2595. doi:10.1016/j.celrep.2016.11.031

17. Pietras EM, Mirantes-Barbeito C, Fong S, et al. Chronic interleukin-1 exposure drives haematopoietic stem cells towards precocious myeloid differentiation at the expense of self-renewal. Nat Cell Biol. 2016;18(6):607–618. doi:10.1038/ncb3346

18. WHO. World Malaria Report 2023.; 2023:287.

19. Coban C, Lee MSJ, Ishii KJ. Tissue-specific immunopathology during malaria infection. Nat Rev Immunol. 2018;18(4): 266–278. doi:10.1038/nri.2017.138

20. Balaji SN, Deshmukh R, Trivedi V. Severe malaria: Biology, clinical manifestation, pathogenesis and consequences. J Vector Borne Dis. 2020;57(1):1–13. doi:10.4103/0972-9062.308793

21. de Sousa LP, de Almeida RF, Ribeiro-Gomes FL, et al. Long-term effect of uncomplicated Plasmodium berghei ANKA malaria on memory and anxiety-like behaviour in C57BL/6 mice. Parasit Vectors. 2018;11(1):191. doi:10.1186/s13071-018-2778-8

22. Lee MSJ, Maruyama K, Fujita Y, et al. Plasmodium products persist in the bone marrow and promote chronic bone loss. Sci Immunol. 2017;2(12):eaam8093. doi:10.1126/sciimmunol.aam8093

23. Alfred Mavondo G, Nkazimulo Mkhwanazi B, Louis Mzingwane M, et al. Malarial Inflammation-Driven Pathophysiology and Its Attenuation by Triterpene Phytotherapeutics. In: Antonio Bastidas Pacheco G, Ali Kamboh A, eds. Parasitology and Microbiology Research. IntechOpen; 2020. doi:10.5772/intechopen.88731

24. Belyaev NN, Brown DE, Diaz AIG, et al. Induction of an IL7-R+c-Kithi myelolymphoid progenitor critically dependent on IFN-γ signaling during acute malaria. Nat Immunol. 2010;11(6):477–485. doi:10.1038/ni.1869

25. Haltalli MLR, Watcham S, Wilson NK, et al. Manipulating niche composition limits damage to haematopoietic stem cells during Plasmodium infection. Nat Cell Biol. 2020;22(12):1399–1410. doi:10.1038/s41556-020-00601-w

26. Spence PJ, Jarra W, Lévy P, Nahrendorf W, Langhorne J. Mosquito transmission of the rodent malaria parasite Plasmodium chabaudi. Malar J. 2012;11(1):407. doi:10.1186/1475-2875-11-407

27. Chen SY, Feng Z, Yi X. A general introduction to adjustment for multiple comparisons. J Thorac Dis. 2017;9(6):1725–1729. doi:10.21037/jtd.2017.05.34

28. Spence PJ, Jarra W, Lévy P, et al. Vector transmission regulates immune control of Plasmodium virulence. Nature. 2013;498(7453):228-231. doi:10.1038/nature12231

29. Pérez-Mazliah D, Ng DHL, Freitas do Rosário AP, et al. Disruption of IL-21 signaling affects T cell-B cell interactions and abrogates protective humoral immunity to malaria. PLoS Pathog. 2015;11(3):e1004715. doi:10.1371/journal.ppat.1004715

30. Achtman AH, Stephens R, Cadman ET, Harrison V, Langhorne J. Malaria-specific antibody responses and parasite persistence after infection of mice with Plasmodium chabaudi chabaudi. Parasite Immunol. 2007;29(9):435–444. doi:10.1111/j.1365-3024.2007.00960.x

31. Belyaev NN, Biró J, Langhorne J, Potocnik AJ. Extramedullary Myelopoiesis in Malaria Depends on Mobilization of Myeloid-Restricted Progenitors by IFN-γ Induced Chemokines. Stevenson MM, ed. PLoS Pathog. 2013;9(6):e1003406. doi:10.1371/journal.ppat.1003406

32. Pietras EM, Reynaud D, Kang YA, et al. Functionally Distinct Subsets of Lineage-Biased Multipotent Progenitors Control Blood Production in Normal and Regenerative Conditions. Cell Stem Cell. 2015;17(1):35–46. doi:10.1016/j.stem.2015.05.003

33. Sommerkamp P, Romero-Mulero MC, Narr A, et al. Mouse multipotent progenitor 5 cells are located at the interphase between hematopoietic stem and progenitor cells. Blood. 2021;137(23):3218–3224. doi:10.1182/blood.2020007876

34. Akinduro O, Weber TS, Ang H, et al. Proliferation dynamics of acute myeloid leukaemia and haematopoietic progenitors competing for bone marrow space. Nat Commun. 2018;9(1):519. doi:10.1038/s41467-017-02376-5

35. Takizawa H, Boettcher S, Manz MG. Demand-adapted regulation of early hematopoiesis in infection and inflammation. Blood. 2012;119(13):2991–3002. doi:10.1182/blood-2011-12-380113

36. Challen GA, Boles N, Lin KK, Goodell MA. Mouse Hematopoietic Stem Cell Identification And Analysis. Cytom Part J Int Soc Anal Cytol. 2009;75(1):14–24. doi:10.1002/cyto.a.20674

37. Mistry JJ, Hellmich C, Moore JA, et al. Free fatty-acid transport via CD36 drives β-oxidation-mediated hematopoietic stem cell response to infection. Nat Commun. 2021;12:7130. doi:10.1038/s41467-021-27460-9

38. Nagai Y, Garrett KP, Ohta S, et al. Toll-like receptors on hematopoietic progenitor cells stimulate innate immune system replenishment. Immunity. 2006;24(6):801–812. doi:10.1016/j.immuni.2006.04.008

39. Isringhausen S, Mun Y, Kovtonyuk L, et al. Chronic viral infections persistently alter marrow stroma and impair hematopoietic stem cell fitness. J Exp Med. 2021;218(12):e20192070. doi:10.1084/jem.20192070

40. de Bruin AM, Voermans C, Nolte MA. Impact of interferon-γ on hematopoiesis. Blood. 2014;124(16):2479–2486. doi:10.1182/blood-2014-04-568451

41. Furusawa J ichi, Mizoguchi I, Chiba Y, et al. Promotion of Expansion and Differentiation of Hematopoietic Stem Cells by Interleukin-27 into Myeloid Progenitors to Control Infection in Emergency Myelopoiesis. Stevenson MM, ed. PLOS Pathog. 2016;12(3):e1005507. doi:10.1371/journal.ppat.1005507

42. Cunnington AJ, Njie M, Correa S, Takem EN, Riley EM, Walther M. Prolonged neutrophil dysfunction after Plasmodium falciparum malaria is related to hemolysis and heme oxygenase-1 induction. J Immunol Baltim Md 1950. 2012;189(11):5336-5346. doi:10.4049/jimmunol.1201028

43. Mooney JP, DonVito SM, Jahateh M, et al. ‘Bouncing Back’ From Subclinical Malaria: Inflammation and Erythrocytosis After Resolution of P. falciparum Infection in Gambian Children. Front Immunol. 2022;13. doi:10.3389/fimmu.2022.780525

44. Pronk CJH, Rossi DJ, Månsson R, et al. Elucidation of the Phenotypic, Functional, and Molecular Topography of a Myeloerythroid Progenitor Cell Hierarchy. Cell Stem Cell. 2007;1(4):428–442. doi:10.1016/j.stem.2007.07.005

45. Wu Q, Zhang J, Kumar S, et al. Resilient anatomy and local plasticity of naive and stress haematopoiesis. Nature. 2024;627(8005):839-846. doi:10.1038/s41586-024-07186-6

46. Hérault A, Binnewies M, Leong S, et al. Myeloid progenitor cluster formation drives emergency and leukaemic myelopoiesis. Nature. 2017;544(7648):53–58. doi:10.1038/nature21693

47. Walter D, Lier A, Geiselhart A, et al. Exit from dormancy provokes DNA-damage-induced attrition in haematopoietic stem cells. Nature. 2015;520(7548):549–552. doi:10.1038/nature14131

48. Cabezas-Wallscheid N, Klimmeck D, Hansson J, et al. Identification of Regulatory Networks in HSCs and Their Immediate Progeny via Integrated Proteome, Transcriptome, and DNA Methylome Analysis. Cell Stem Cell. 2014;15(4):507–522. doi:10.1016/j.stem.2014.07.005

49. Yamashita M, Passegué E. TNF-α Coordinates Hematopoietic Stem Cell Survival and Myeloid Regeneration. Cell Stem Cell. 2019;25(3):357–372.e7. doi:10.1016/j.stem.2019.05.019

50. Kang YA, Paik H, Zhang SY, et al. Secretory MPP3 reinforce myeloid differentiation trajectory and amplify myeloid cell production. J Exp Med. 2023;220(8):e20230088. doi:10.1084/jem.20230088

51. Johnson NB, Posluszny JA, He LK, et al. Perturbed MafB/GATA1 axis after burn trauma bares the potential mechanism for immune suppression and anemia of critical illness. J Leukoc Biol. 2016;100(4):725–736. doi:10.1189/jlb.1A0815-377R

52. Lin F ching, Karwan M, Saleh B, et al. IFN-γ causes aplastic anemia by altering hematopoietic stem/progenitor cell composition and disrupting lineage differentiation. Blood. 2014;124(25):3699–3708. doi:10.1182/blood-2014-01-549527

53. Shi X, Wei S, Simms KJ, Cumpston DN, Ewing TJ, Zhang P. Sonic Hedgehog Signaling Regulates Hematopoietic Stem/Progenitor Cell Activation during the Granulopoietic Response to Systemic Bacterial Infection. Front Immunol. 2018;9. doi:10.3389/fimmu.2018.00349

54. Martinez A, Bono C, Megías J, Yáñez A, Gozalbo D, Gil ML. Systemic Candidiasis and TLR2 Agonist Exposure Impact the Antifungal Response of Hematopoietic Stem and Progenitor Cells. Front Cell Infect Microbiol. 2018;8. doi:10.3389/fcimb.2018.00309

55. Hümmer J, Kraus S, Brändle K, Lee-Thedieck C. Nitric Oxide in the Control of the in vitro Proliferation and Differentiation of Human Hematopoietic Stem and Progenitor Cells. Front Cell Dev Biol. 2021;8:610369. doi:10.3389/fcell.2020.610369

56. Lee MSJ, Maruyama K, Fujita Y, et al. Plasmodium products persist in the bone marrow and promote chronic bone loss. Sci Immunol. 2017;2(12):eaam8093. doi:10.1126/sciimmunol.aam8093

57. Torre D, Speranza F, Giola M, Matteelli A, Tambini R, Biondi G. Role of Th1 and Th2 Cytokines in Immune Response to Uncomplicated Plasmodium falciparum Malaria. Clin Vaccine Immunol. 2002;9(2):348–351. doi:10.1128/CDLI.9.2.348-351.2002

58. Luty AJF, Lell B, Schmidt-Ott R, et al. Interferon-γ Responses Are Associated with Resistance to Reinfection with Plasmodium falciparum in Young African Children. J Infect Dis. 1999;179(4):980–988. doi:10.1086/314689

59. Takem EN, Roca A, Cunnington A. The association between malaria and non-typhoid Salmonella bacteraemia in children in sub-Saharan Africa: a literature review. Malar J. 2014;13:400. doi:10.1186/1475-2875-13-400

