## Supplemental figures for "Differential responses and recovery dynamics of HSPC populations following *Plasmodium chabaudi* infection"

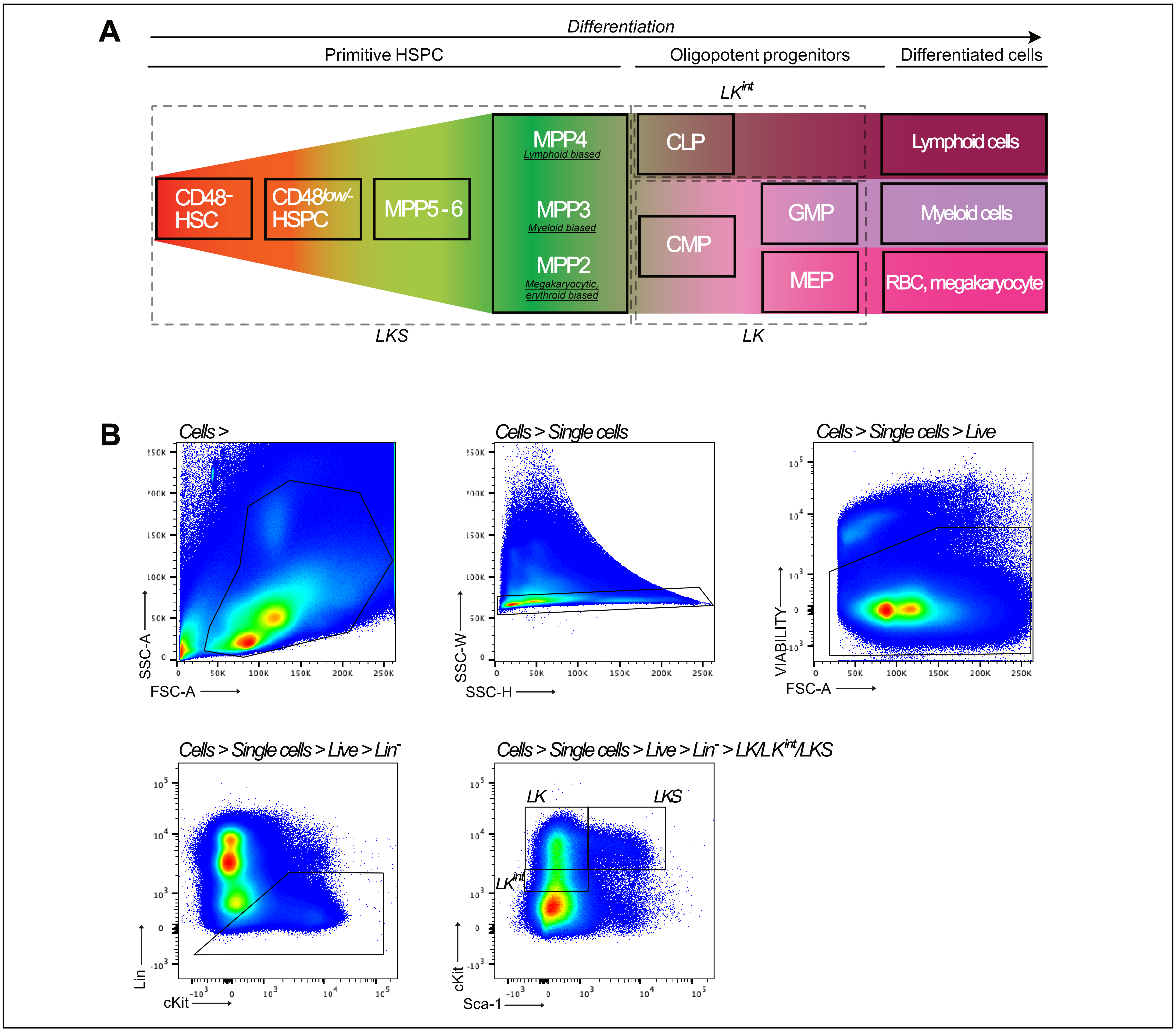


**Figure S1. Hematopoietic populations investigated in response to P. chabaudi infection**

**A.** Schematic representation summarizing the hierarchy of the different hematopoietic populations and the nomenclature used in this study. LKS cells include primitive HSPC, thus HSCs and MPPs. Oligopotent progenitors are divided into LK and $LK^{int}$ cells. LK cells include CMP upstream of GMP and MEP populations, and $LK^{int}$ cells include CLPs. **B.** Flow cytometry plots showing the gating strategy used from all events to LK/LKS populations. The gating strategies for further subpopulations are shown in other figures.

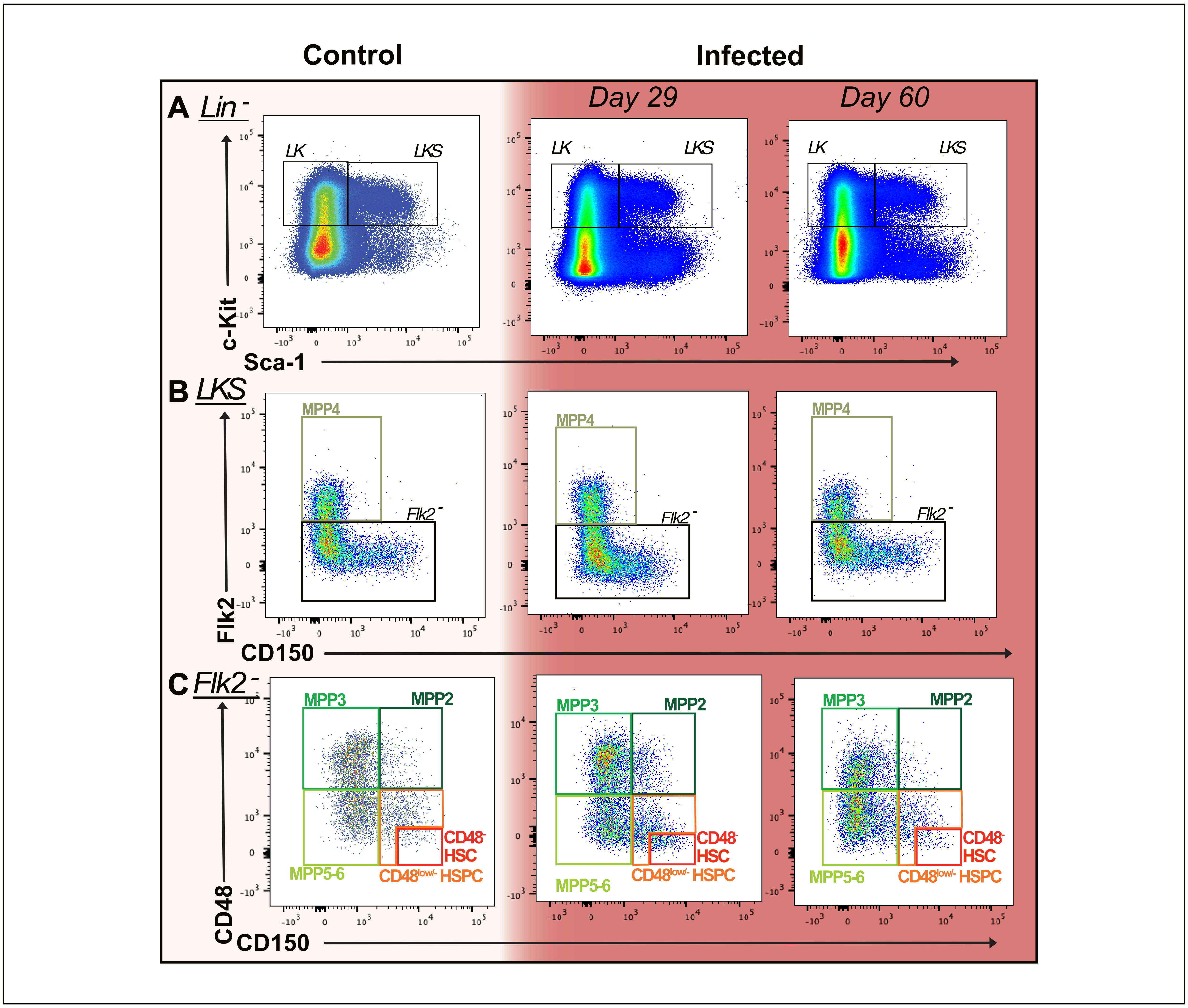


**Figure S2: Primitive hematopoietic populations during the recovery phase of *P. chabaudi* infection.**

**A-E**. Representative flow cytometry plots showing LK and LKS (**A**), MPP4(**B**), MPP2, MPP3, MPP5-6, CD48- HSC and CD48low/- HSPC (**C**) populations in controls and infected mice at day 29 and day 60 p.i.

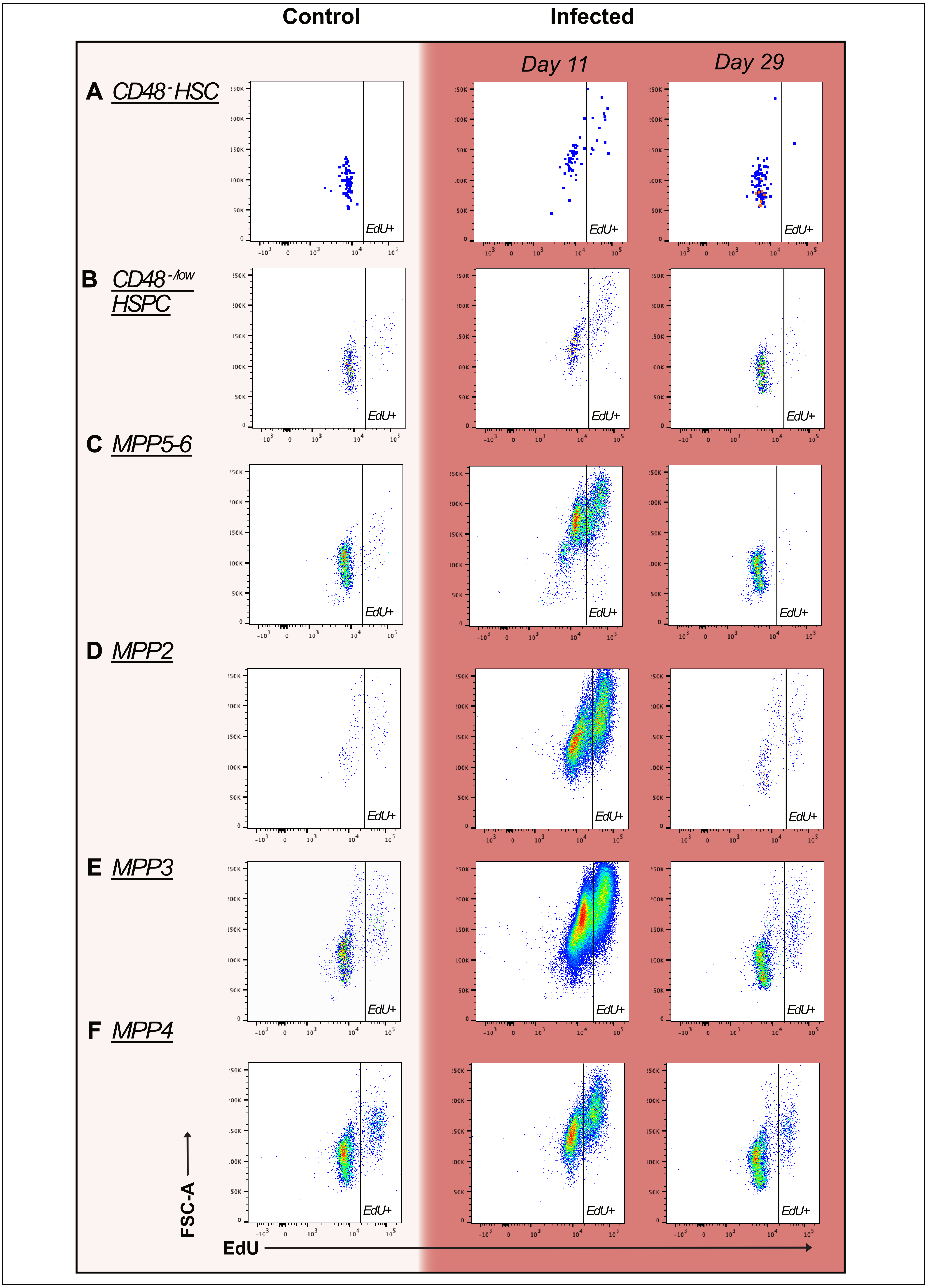


**Figure S3: Proliferation of primitive hematopoietic populations during *P. chabaudi* infection.**

**A-E**. Representative flow cytometry plots showing cells in the S-phase of the cell cycle (EdU$^+$) within CD48$^-$ HSC (**A**), CD48$^{low/-}$ HSPC (**B**), MPP5-6(**C**), MPP2 (**D**), MPP3 (**E**) and MPP4 (**F**) populations in controls and infected mice at day 11 and day 29 p.i.

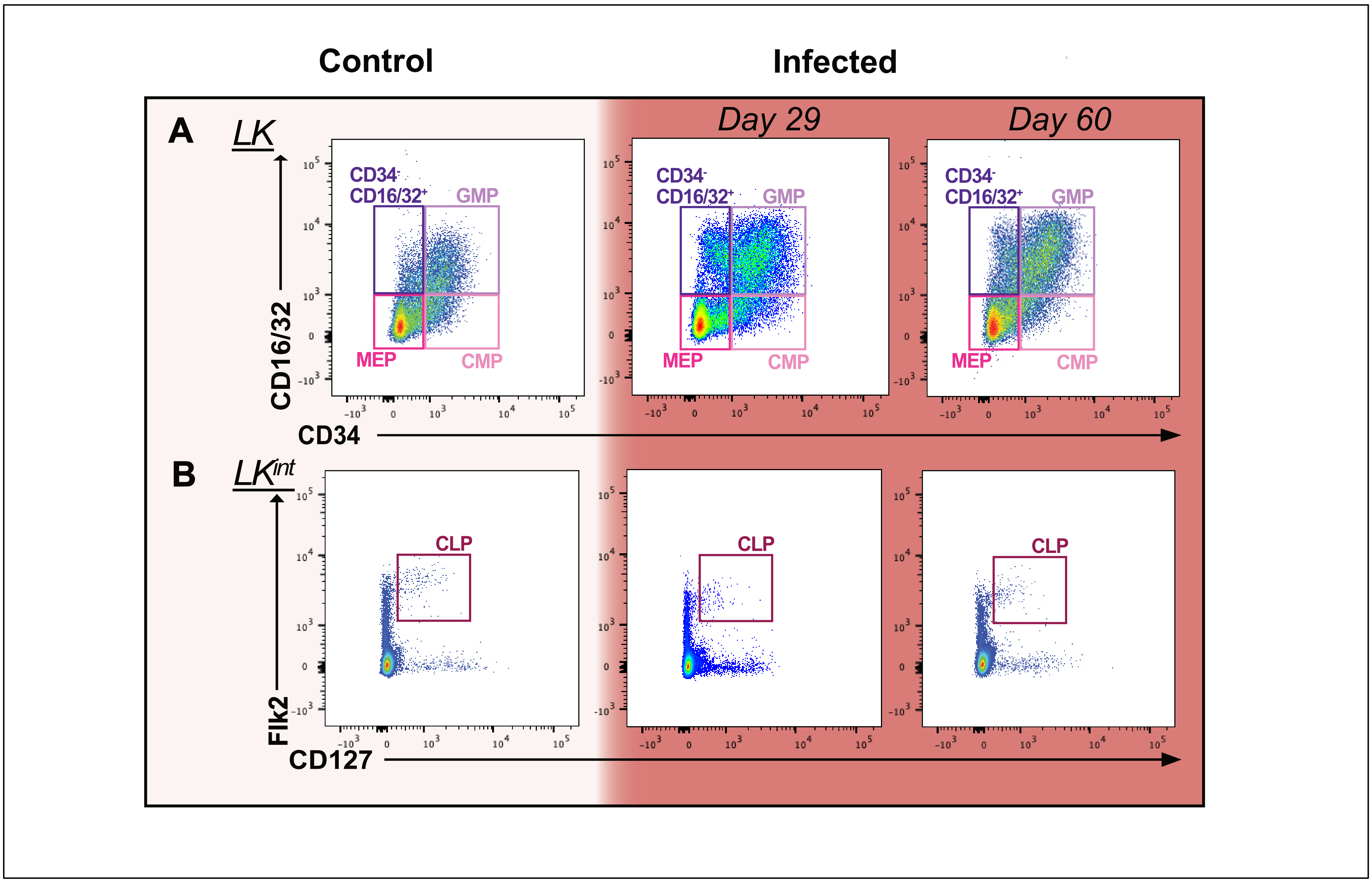


**Figure S4: Hematopoietic populations during the recovery phase of *P. chabaudi* infection.**

**A-E**. Representative flow cytometry plots showing (**A**) CD34-CD16/32+, GMP, CMP and MEP populations within the LK parent population and (**B**) CLP cells within the parent LK$^{int}$ population in controls and infected mice at day 29 and day 60 p.i.

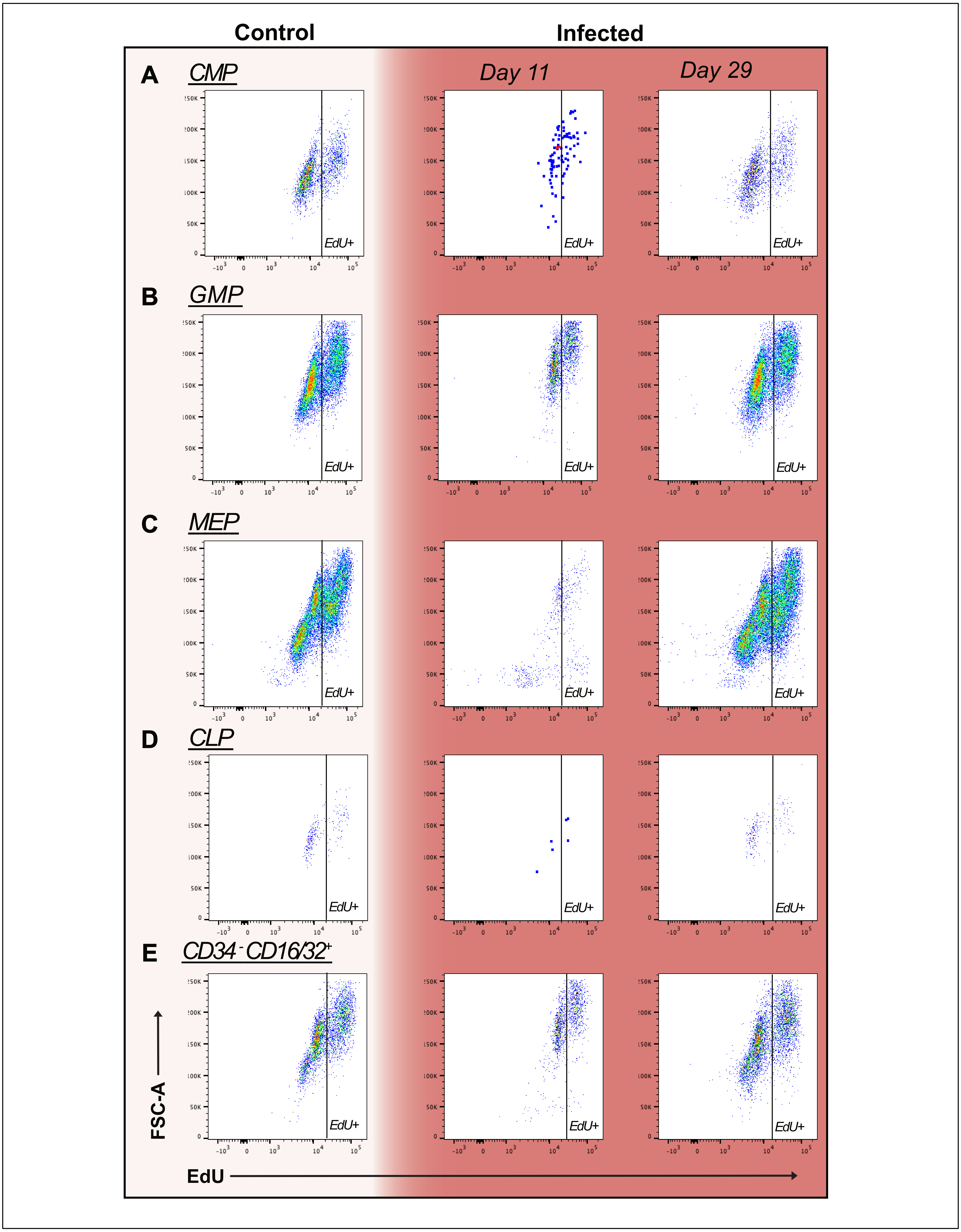


**Figure S5: Proliferation of mature progenitor populations during *P. chabaudi* infection.**

**A-E**. Representative flow cytometry plots showing in the S-phase of the cell cycle (EdU$^+$) within CMP (**A**), GMP (**B**), MEP (**C**), CLP (**D**) and CD34$^-$CD16/32$^+$ (**E**) populations in controls and infected mice at day 11 and day 29 p.i.

| Antibody | Clone | Manufacturer | Cat. no | Dilution |
|---|---|---|---|---|
| FITC anti-mouse CD4 | RM4-5 | Biolegend | 100510 | 1/400 |
| FITC anti-mouse CD5 | 53-7.3 | Biolegend | 100606 | 1/100 |
| FITC anti-mouse CD8a | 53-6.7 | Biolegend | 100706 | 1/200 |
| FITC anti-mouse B220 | RA3-6B2 | Biolegend | 103206 | 1/200 |
| FITC anti-mouse Ter119 | TER-119 | Biolegend | 116206 | 1/100 |
| FITC anti-mouse Ly-6G/Ly-6C (Gr-1) | RB6-8C5 | Biolegend | 108406 | 1/400 |
| APC/Cyanine7 anti-mouse CD117 (c-kit) | 2B8 | Biolegend | 105826 | 1/200 |
| PerCP/Cyanine5.5 anti-mouse Ly-6A/E (Sca-1) | D7 | Biolegend | 108124 | 1/300 |
| Brilliant Violet 650 anti-mouse CD150 | TC15-12F12.2 | Biolegend | 115931 | 1/200 |
| PE/Cyanine7 anti-mouse CD48 | HM48-1 | Biolegend | 103424 | 1/200 |
| PE anti-mouse CD127 | A7R34 | Invitrogen | 12-1271-82 | 1/100 |
| FITC anti-mouse CD34 | RAM34 | Invitrogen | 11-0341-81 | 1/50 |
| APC anti-mouse CD135 | A2F10 | Biolegend | 135310 | 1/200 |

**Table S1. Details of antibodies used in this study.**
